# Specialized signaling centers direct cell fate and spatial organization in a limb organoid model

**DOI:** 10.1101/2024.07.02.601324

**Authors:** Evangelia Skoufa, Jixing Zhong, Oliver Kahre, Kelly Hu, Georgios Tsissios, Louise Carrau, Antonio Herrera, Albert Dominguez Mantes, Alejandro Castilla-Ibeas, Hwanseok Jang, Martin Weigert, Gioele La Manno, Matthias Lutolf, Marian Ros, Can Aztekin

**Affiliations:** School of Life Sciences, Swiss Federal Institute of Technology Lausanne (EPFL), Lausanne 1015, Switzerland; Department of Cellular and Molecular Signalling, Instituto de Biotecnología y Biomedicina de Cantabria (IBBTEC), CSIC-SODERCAN-University of Cantabria), Santander, Spain; Institute of Human Biology (IHB), Roche Pharma Research and Early Development, Roche Innovation Center, Basel 4058, Switzerland

## Abstract

Specialized signaling centers orchestrate robust development and regeneration. Limb morphogenesis, for instance, requires interactions between the mesoderm and the signaling center apical-ectodermal ridge (AER), whose properties and role in cell fate decisions have remained challenging to dissect. To tackle this, we developed mouse embryonic stem cells (mESCs)-based heterogeneous cultures and a limb organoid model, termed budoids, comprising cells with AER, surface ectoderm, and mesoderm properties. mESCs were first induced into heterogeneous cultures that self-organized into domes in 2D. Aggregating these cultures resulted in formation of limb bud-like structures in 3D, exhibiting chondrogenesis-based symmetry breaking and elongation. Using our organoids and quantitative in situ expression profiling, we uncovered that AER-like cells support nearby limb mesoderm and fibroblast identities while enhancing tissue polarization that permits distant cartilage formation. Together, our findings provide a powerful model to study aspects of limb morphogenesis, and reveal the ability of signaling center AER cells to concurrently modulate cell fate and spatial organization.

## Introduction

Specialized signaling centers are unique cell populations, defined by their diverse morphogen secretion capabilities, that transiently form and orchestrate morphogenesis during development and regeneration^1^. A notable example is found in vertebrate limbs, where surface ectoderm cells differentiate into the signaling center apical-ectodermal ridge (AER) cells. The AER then establishes multiple morphogen gradients (including FGFs, BMPs, WNTs, TGFBs, and DELTAs ^2,3^) simultaneously to influence the growth and differentiation of underlying multipotent limb bud mesoderm. It has been postulated that mesodermal cells leaving the AER morphogen gradient zone differentiate into chondrocytes and fibroblasts^2^. In parallel, signals from the non-AER surface ectoderm have been suggested to suppress chondrocyte differentiation^4^. Although it is crucial to understand how these intricate cell-cell interactions are orchestrated to shape a developing limb, studies on signaling centers and limb morphogenesis are severely restricted by *in vivo* experimentation, which is constrained to tissue or gene-level examinations. The lack of viable cellular platforms poses a significant challenge to using large-scale and quantitative measurements to study specialized signaling centers and their cell-cell interactions guiding limb development and regeneration.

Due to their well-controlled make-up, the ease of observation and perturbation, and scalability, organoids have become powerful models to yield new insights into the principles of morphogenesis^5^. While many organoids and developmental phenomena rely on single-lineage self-organization^6^, limbs require multi-lineage orchestration between ectoderm and mesoderm^7,8^. However, simplified limb models have almost exclusively focused on the limb bud mesoderm^9–13^, and the exploration of the ectoderm, including the AER, has been hampered by the unavailability of a suitable model^14^. A recent 3D limb bud model incorporated mesodermal and ectodermal cells, but it required manual dissection for organoid generation, fell short in emulating robust morphogenesis, and lacked comprehensive single-cell and functional characterization^13^. Therefore, we developed and characterized a highly efficient *in vitro* limb organoid system from mouse embryonic stem cells (mESCs) to dissect the self-organization propensities of distinct cell types and the specific roles of signaling center AER cells.

## Results

### Generation and characterization of stem-cell-derived heterogenous cultures resembling limb bud cell types

We initially aimed to only generate signaling center AER cells from mESCs (Figure 1A-B). Adapting a published protocol^15^ (via SB431542 and BMP4) to induce surface ectoderm-like cells—a precursor population to AER—yielded homogenous epithelial cells (Figure S1A-C). Subsequent activation of pathways required for AER induction, via BMP4^16^, FGF10^17^, and Wnt^18^ agonist Chiron (hereafter, BFC) treatment, upregulated well-established AER genes such as *Fgf8, Msx2, Wnt5a*^2^ (Figure S1E). Putative AER-like cells were detected as clusters based on *Fgf8* and TP63 staining on day 7 (Figure S1F). Generating and using a *Fgf8: tdTomato* reporter knock-in line (Figure S1G-H), we further found clusters of *Fgf8:tdTomato+*/TP63+ cells (Figure 1C and Figure S1I).

**Figure 1:**
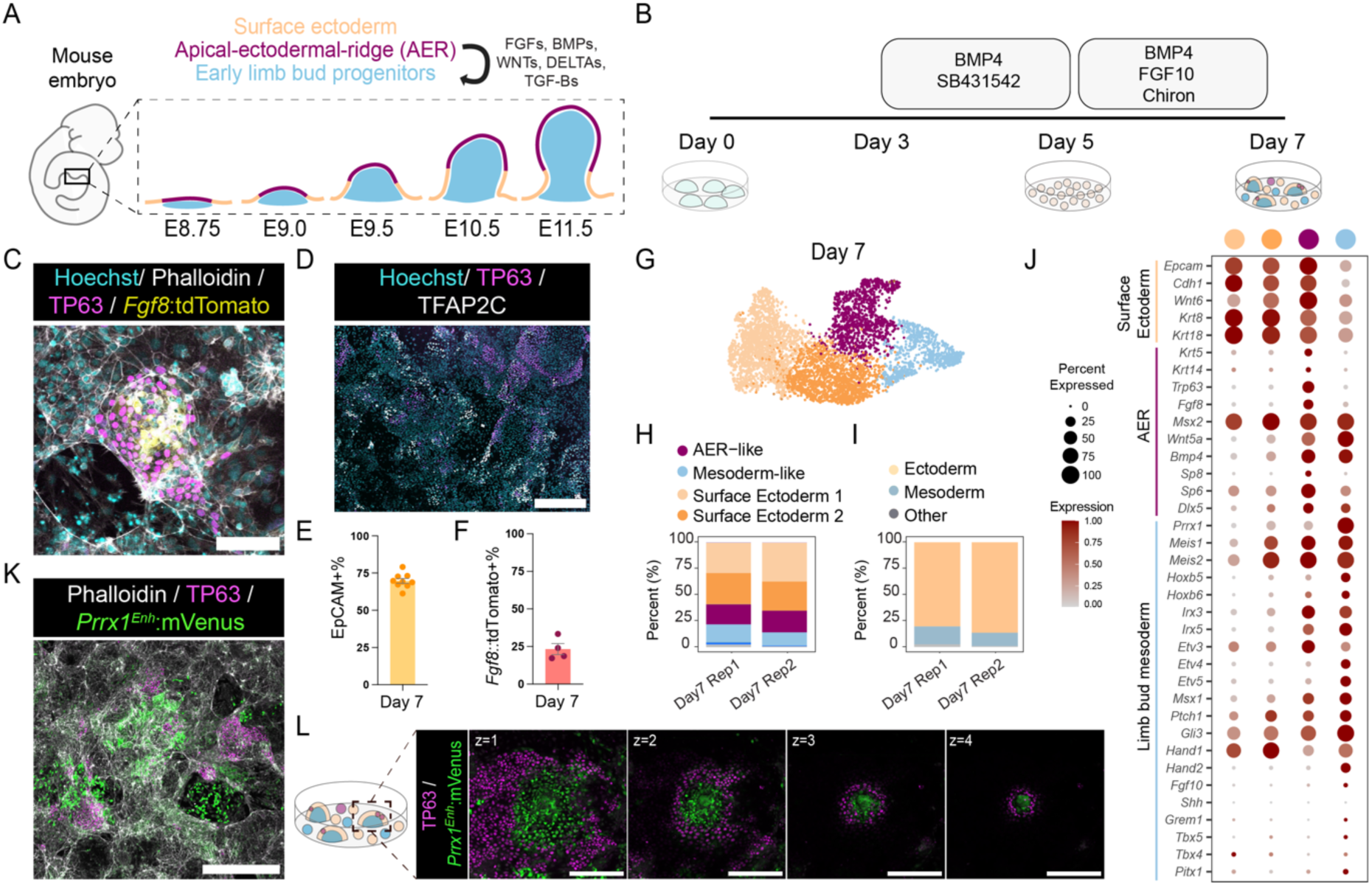
Induced heterogeneous cultures contain cells with signaling center AER, surface ectoderm, and mesoderm properties that self-organize into domes. (A) Schematic describing the embryonic stages of mouse limb bud formation and the main cell populations involved in limb morphogenesis. (B) Schematic of the protocol to generate cells with AER, surface ectoderm, and mesoderm properties from mouse embryonic stem cells (mESCs), with treatments indicated at each time point. (C) Representative confocal image of generated AER-like cells in Day 7 cultures. Cyan: Hoechst, White: Phalloidin, Magenta: TP63, Yellow: *Fgf8:tdTomato*. Scale bar =100 μm. (D) Representative confocal image of Day 7 heterogeneous cultures stained for immature surface ectoderm TFAP2C and basal epithelial stem cell marker TP63. Cyan: Hoechst, Magenta: TP63, White: TFAP2C. Scale bar = 400 μm. (E) Bar plot showing flow cytometry quantification of epithelial cell marker EpCAM positive cells in Day 7 cultures. Each dot represents one biological replicate. N=9. (F) Bar plot showing flow cytometry quantification of *Fgf8:tdTomato* positive cells in Day 7 cultures. Each dot represents one biological replicate. N=4. (G) UMAP representation of scRNA-Seq data of day 7 cultures displaying main cell clusters. For color coding see panel H (H-I) Chart plots showing the scRNA-Seq cluster-based proportions of (H) generated cell types, with color coding for AER-like, mesoderm-like, and surface ectoderm, and (I) lineages on day 7 of the protocol. For full information, please see Figure S2E. Rep denotes replicate. (J) Dot plot showing surface ectoderm, AER, and limb bud mesoderm marker gene expression in the scRNA-Seq clusters. The size of the dot represents the percentage of cells expressing each marker. (K) Representative confocal image of day 7 heterogeneous cultures generated from the *Prrx1^Enh^:mVenus* reporter mESC line. White: Phalloidin, Magenta: TP63, Green: *Prrx1^Enh^:mVenus*. Scale bar = 500 μm. (L) Representative sequential Z-stack confocal images of dome formation in Day 7 cultures. Magenta: TP63, Green: *Prrx1^Enh^:mVenus*. Images from left to right show the dome from the bottom to the top, denoted by z numbers. Scale bar = 200 μm.

However, not all generated cells expressed these markers or the more immature surface ectoderm marker TFAP2C (Figure 1D). Furthermore, flow cytometry detected 69.7(±1.6)% epithelial-cell-marker EpCAM+ (Figure 1E and Figure S1J), and 23.3(±3.6)% *Fgf8:tdTomato*+ cells (Figure 1F and Figure S1K). The cultures also contained cells with fibroblast morphology (Figure S1D) and showed mesoderm-related *Prrx1* expression (Figure S1L), indicating the simultaneous formation of epithelial and mesenchymal populations.

### Single cell characterization of heterogenous cultures and their similarity to *in vivo* limb buds

We reasoned that the generated AER-like cells were altering the local environment due to their ligand secretion capacity, resulting in the induction of heterogeneous cell types within the culture. To determine the cellular composition of our cultures, we performed time-course (0, 3, 5, and 7 days) single-cell mRNA-sequencing (scRNA-Seq), and clustering-based cell type annotation (Figure S2A-F). Days 0 to 3 of culture showed the expected transition from pluripotency states and the formation of neuroectodermal cells (Figure S2A-F). Day 5 contained clusters expressing non-neural ectoderm markers (e.g., *Krt8*, and *Krt18*) (Figure S2A-F), and some of these cells had an epithelial-to-mesenchymal transition signature (e.g., *Snai1/2, Zeb1/2*) (Figure S2G-H), which might be associated with the induced mesodermal cells and fibroblasts following the BFC treatment.

Day 7 samples induced by the BFC treatment were mainly comprised of cells with AER, surface ectoderm, or mesoderm features (Figure 1G-I, Figure S2A-F). The AER-like cluster was detected by associated transcription factors (e.g., *Sp6, Sp8*^19^), and various signaling ligands (e.g., *Fgf8, Bmp4, Wnt5a*). Within our day 7 cultures, this cluster had *Trp63* expression as a unique marker (Figure 1J, Figure S2D). Although some of these cells had high *Fgf8* expression (Figure 1J), they largely lacked other AER-FGFs (*Fgf4*, *Fgf9, Fgf17)* (Figure S3A), suggesting they might represent an initial/early forming AER state. Moreover, the generated AER-like cells were mainly in the G1 cell cycle phase (Figure S2F), consistent with the known mitotic inactivity of the AER^20,21^. Compared to publicly available limb datasets from E9.5-E12.5^22^, these cells had the highest similarity to *in vivo* E9.5 AER (Figure S3B). Meanwhile, most BFC-induced cells showed surface-ectoderm markers (e.g., *Krt8, Wnt3, Wnt6*) (Figure 1J, Figure S2D). None of these cells showed expression of other relevant ectoderm or neural crest markers (Figure S3C).

Surprisingly, the day 7 mesoderm cluster had lateral plate and early limb bud mesoderm marker expressions (e.g., *Hand1, Hand2,* and *Prrx1*) (Figure 1J, Figure S2D), while lacking marker genes for other mesodermal populations (e.g., cranial mesoderm) (Figure S3E). Moreover, they did not have high expression of *Shh* or *Grem1*, nor hindlimb or forelimb-associated genes (e.g., *Tbx4/5*) (Figure 1J, Figure S2D). Unlike *in vivo* highly proliferative limb bud mesoderm^23^, they were mostly in G1 phase (Figure S2F). To further examine their identity, we compared our data to reference E9.5-E12.5 mouse limb datasets^22^. Correlation analysis showed a putative matching across previously annotated^22^ limb bud mesodermal populations, albeit the absolute values of correlations were low (Figure S3B, D). To further test if cultures contain limb bud mesoderm-like cells, we generated a mESC reporter line containing the *Prrx1* enhancer, traditionally used to label limb bud mesoderm with high precision^24,25^. Using this line, we identified that our protocol produces *Prrx1*^enh^+ fibroblast-like and more circular-looking cells (Figure 1K, Figure S4A-B).

In summary, our protocol induced a heterogeneous population composed of cells that show significant similarities to AER, surface ectoderm, and, to some degree, early lateral plate mesoderm and limb bud mesoderm identities.

### Heterogenous cultures self-organize into domes in 2D and budoids in 3D

Next, we examined the spatial organization of the generated AER-like and mesodermal cells in our 2D cultures. Remarkably, we found that the generated cells self-organized into 3D “domes” with TP63+ cells covering the outer layers of the domes and TP63+/*Fgf8*:*tdTomato*+ AER-like cells at the tip or outer bottom layers (Figure 1L, Figure S4C, E, F). Moreover, the core of the domes was populated by *Prrx1^enh^*+ cells (Figure 1L), or LEF1+/TP63-/TFAP2C-cells with smaller nuclei (Figure S4D-F), corresponding to the mesodermal cluster in the scRNA-Seq dataset (Figure S4D). Together, these experiments suggested self-organization via AER-like and mesoderm interactions, generating domes in 2D cultures that extend into 3D and resemble limb buds.

We subsequently asked if our generated cells can also self-organize in 3D as free-floating cultures (Figure 2A). Dissociating and seeding cultures in U-bottom low-attachment plates resulted in cells forming single spheric aggregates. Moreover, 5.08(±2.9)% of these aggregates displayed limited symmetry breaking and elongation (Figure 2B, Figure S5A-B). Since it is known that close-range ectoderm-mesoderm interactions are required for limb morphogenesis^7,8^, we reasoned that the low degree of symmetry breaking might be due to insufficient ectodermal signals. Indeed, a brief treatment of ectodermal signals FGF8b and WNT3A during aggregation significantly increased the symmetry break efficiency to an average of 80.7(±5.9)% with a greater elongation phenotype (Fig 2B-D, Figure S5C-E). As our 3D structures were generated from cells with limb development characteristics and showed morphological characteristics of limb buds, we termed them *budoids*.

**Figure 2:**
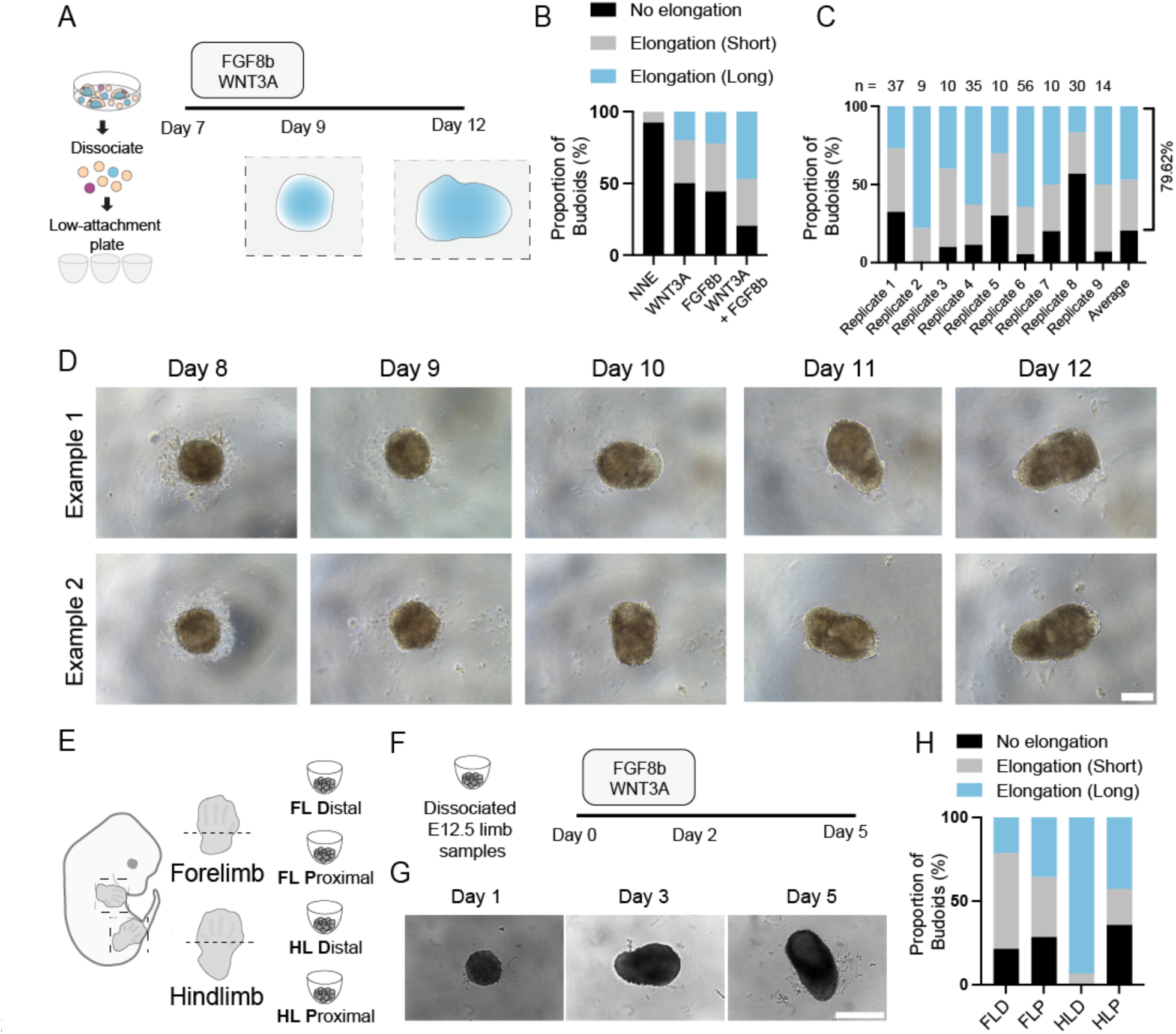
Stem-cell-derived heterogeneous cultures and *in vivo* limb cells can form budoids. (A) Schematic illustration of the budoid generation protocol. Treatments and morphological changes to cell aggregates at designated time points from Day 7 to Day 12 are outlined. (B) Bar graph displaying the proportion of budoids with no elongation, elongation (short), or elongation (long) after brief treatment with 300 ng/ml WNT3A, 450 ng/ml FGF8b, or a combination of both compared to control (NNE, non-neural ectoderm media). NNE: n= 39 budoids, N=3 : 300 ng/ml WNT3A n=10, N=1; 450 ng/ml FGF8b n=9, N=1: 300 ng/ml WNT3A and 450 ng/ml FGF8b, n=72, N=2. (C) Bar graph showing the percentage of budoid structures achieving different elongation phenotypes under brief 300 ng/ml WNT3A and 450 ng/ml FGF8b treatment. The number above each set of bars (n) indicates the number of budoids analyzed within the replicate. Last column shows weighted average. The total number of budoids analyzed was n= 211, N=9 replicates. (D) Sequential brightfield images of examples of budoids from Day 8 to Day 12 of the protocol. Scale bar = 100 μm. (E) Schematic representing dissection of forelimbs and hindlimbs into distal and proximal sections for budoid generation from primary cells. (F) Timeline schematic showing the dissociation of E12.5 limb samples and subsequent treatment with FGF8b and WNT3A at specified days to promote budoids formation and elongation. (G) Representative brightfield images of primary cell-derived hindlimb distal (HLD) budoids at Days 1, 3, and 5 of the protocol. Scale bar = 200 μm (H) Bar graph illustrating the proportion of bud-like structures from forelimb distal (FLD), forelimb proximal (FLP), hindlimb distal (HLD), and hindlimb proximal (HLP) regions, categorized by no elongation, elongation (short), or elongation (long) after 5 days of culture. Total number of budoids analyzed FLD n=14, N=3; FLP n=14, N=3; HLD n=15: N=3; HLP, n=14, N=3.

Our results indicated that our stem cell-derived cells with limb developmental features can self-organize in 3D where they aggregate, subsequently break symmetry and elongate. To further investigate the degree to which our cultures resembled *in vivo* limb bud cell types, we evaluated whether developing limb cells from embryonic mice could similarly self-organize in 3D in our setting. For this, we harvested and dissociated E12.5 mouse distal and proximal limbs (Figure 2E) and exposed these to our protocol. All samples, particularly the hindlimb distal sample, underwent morphological organization similar to those seen in stem cell-derived budoids (Figure 2F-H, Figure S6). Overall, these results demonstrated that our simplified setting can uncover morphogenetic capacity of both stem cell- and *in vivo*-derived limb developmental cells.

### Budoids break symmetry through limb mesoderm-to-chondrocyte differentiation

To unravel cell fate changes during stem cell-derived budoid morphogenesis, we performed scRNA-Seq and clustering analysis on pooled budoid samples before (day 9) and after symmetry breaking (day 12), and combined the generated datasets with our 2D heterogeneous culture data as a starting point (Figure 3A-B, Figure S7).

**Figure 3:**
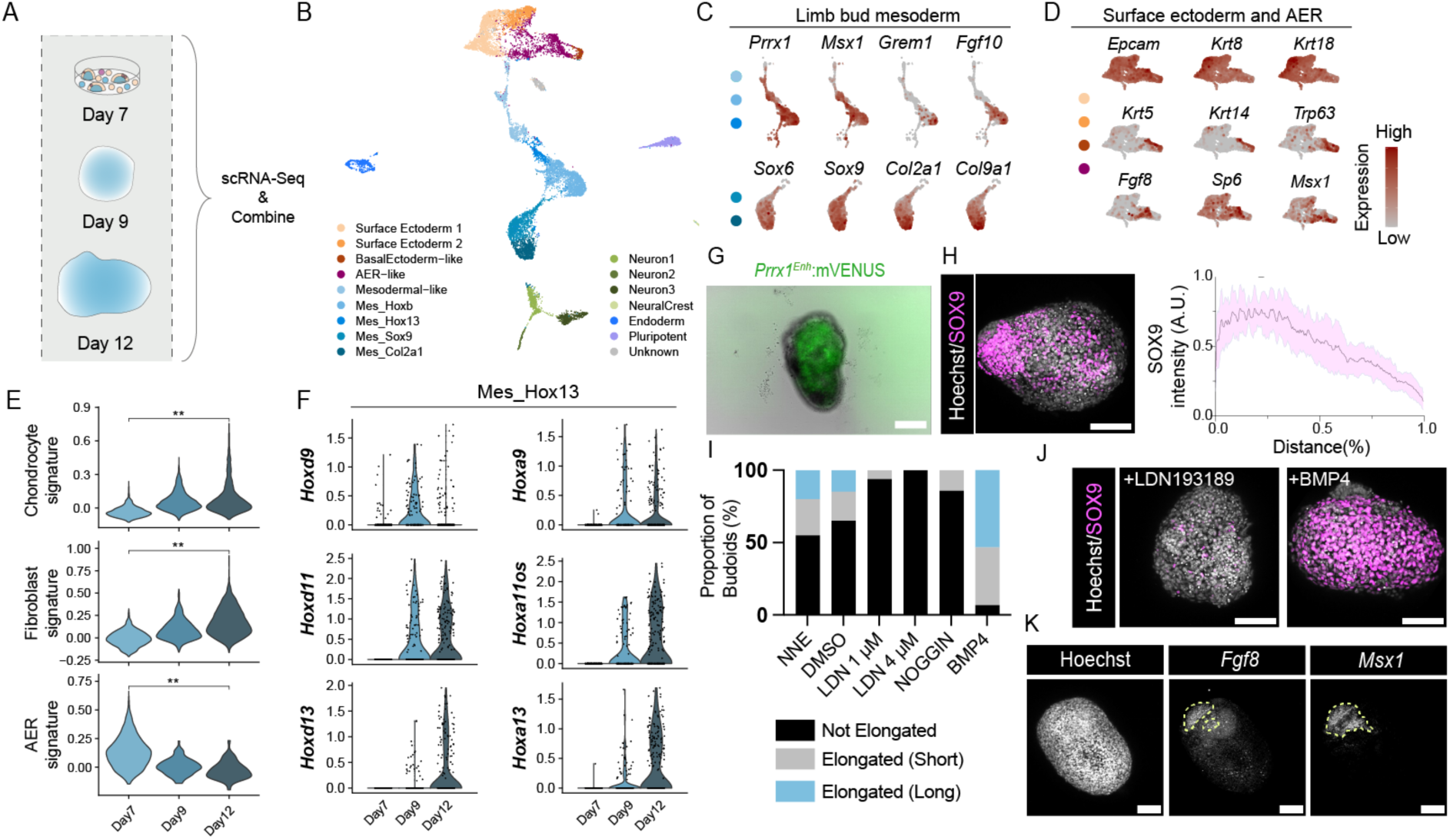
Budoids exhibit limb developmental hallmarks, and BMP-regulated mesoderm to chondrocyte differentiation mediates their symmetry breaking and elongation. (A) Schematic overview of the scRNA-Seq analysis process with samples combined from different time points. (B) UMAP representation of the combined data illustrating the diverse cell populations identified in the scRNA-Seq analysis. (C) Feature plots displaying the expression of key limb bud mesoderm and chondrogenesis genes in mesodermal clusters indicated by the color dots on the left. (D) Feature plots showing the expression of surface ectoderm and AER genes in ectodermal clusters, indicated by the color dots on the left. (E) Violin plots depicting gene set enrichment scores for signatures associated with chondrocyte and fibroblast identities in combined mesodermal clusters and AER cell identity in combined ectodermal clusters, tracked across different days of budoid generation. Two-sided Wilcox test was performed to compare day 7 and day 12 data. ** indicates *p* value < 0.01. (F) Violin plots demonstrating the time course of 5′ *HoxA and HoxD* gene activation in the Mes_Hox13 cluster. Each dot represents one cell. (G) A representative brightfield and fluorescent overlaid image of a day 12 budoid generated using *Prrx1^Enh^:mVenus* line. Green: *Prrx1^Enh^:mVenus*. Scale bar = 100 μm. (H) (Left) A representative max-projection confocal image displaying the polarized expression of SOX9 within a day 13 budoid. Gray: Hoechst, Magenta: SOX9. Scale bar = 100 μm. (Right) Normalized SOX9 intensity across the major axis on day 12 budoids. Total number of analyzed budoids n= 10, N= 4. (I) Bar chart displaying the weighted average of proportion of day 13 budoids exhibiting no elongation, elongation (short), or elongation (long) after treatment with DMSO control, 1 or 4 μM LDN193189, 300 ng/ml recombinant NOGGIN, or 300 ng/ml recombinant BMP4. NNE (no treatment control): n=20, N=2; DMSO: n= 20, N=2; 1 μM LDN193189 (LDN 1 μM): n= 16, N=2; 40 μM LDN193189 (LDN 4 μM): n= 19, N=2; 300 ng/ml recombinant NOGGIN: n= 14, N=2; 300 ng/ml recombinant BMP4: n= 35, N=2. (J) Max-projection confocal images of day 13 budoids with 1 μM LDN193189 or 300 ng/ml BMP4 treatments. Gray: Hoechst, Magenta: SOX9. Scale bar = 100 μm. (K) Confocal image of a budoid showing the polarized expression patterns of *Fgf8* mRNA and *Msx1* mRNA positive cells. Please note that this phenotype is seen in 3/10 budoids. Scale bar = 100 μm.

ScRNA-Seq demonstrated enrichment of mesodermal cells (Figure 3C-G, Figure S7B), with budoids containing cells with multipotent limb bud mesoderm markers (*e.g., Msx1, Grem1, Fgf10*)^26^ (Figure 3C, E-G, Figure S7F) including sequential activation of *Hox* clusters (Figure 3F) and broad significant transcriptome-wide similarity to *in vivo* limb bud mesodermal cells (Figure S8A). We then showed that budoids mainly comprise *Prrx1^Enh^*+ cells (Figure 3G), in agreement with our single-cell analyses. Anterior-posterior patterning-related *Shh* was seen in a few cells (Figure S7F). Mesodermal cells were agnostic for hindlimb (*Tbx4, Pitx1),* or forelimb gene profiles (*Tbx5*) (Figure S7F), with no single cell showing abundant *Tbx4* and *Tbx5* expression simultaneously (Figure S7F, Figure S8B). Consistent with the hypothesis that budoids contain multipotent limb bud mesoderm-like cells, we found budoids to express chondrocytes (*Sox9, Col2a1*), and fibroblast-associated genes (*Col1a1, Col3a1, Dpt*), including tenocytes (*Scx, Lum)* and pericytes (*Rgs5, Acta2, Des)* (Figure S7B).

Critically, post symmetry breaking, scRNA-Seq indicated an increase in chondrocyte populations in budoids (Figure S7B). This enrichment was corroborated by our imaging results where we observed the emergence of a polarized SOX9+ domain (Figure 3H), which was also observed in *in vivo*-derived budoids (Figure S8C). Thus, we hypothesized that budoid symmetry breaking is mediated by chondrogenesis. Indeed, suppression of the well-established chondrogenesis regulator BMP pathway by addition of recombinant NOGGIN or pharmacological inhibition via LDN-193189, impaired symmetry breaking and elongation (Figure 3I), and these budoids had significantly fewer SOX9+ cells (Figure 3J). In contrast, the addition of recombinant BMP4 showed the opposite effect (Figure 3I), with almost all organoids elongated and entirely populated by SOX9+ cells (Figure 3J). Collectively, these results demonstrated that the budoid symmetry break is mediated by limb mesoderm to chondrocyte differentiation, which is regulated by BMP pathway activity.

Substantial ectodermal reduction was evident during budoid formation (Figure S7B), reinforced by the observation that budoids had limited E-CADHERIN+, and TP63+ cells (Figure S8D). We also detected the same phenotype with *in vivo*-derived budoids (Figure S8C), suggesting that our culture conditions will require further optimization for ectodermal cell aggregation and survival. Moreover, the detected ectodermal cells had a lower AER signature compared to day 7 samples (Figure 3E). Nonetheless, a spatially distinct AER marker *Fgf8* was identified as coexisting with regions of *Msx1* expression as has been seen *in vivo* ^27^, albeit this phenotype was only observed in some budoids (Figure 3K). Budoids also had some non-limb lineages (e.g., *Sox17*+ endodermal cells*, Nanog*+ stem cells) (Figure S7B), presumably due to differentiation artifacts that are commonly seen in stem-cell-derived organoids^28^. Collectively, these findings suggest that budoids are organoids, mainly composed of mesodermal cells, that show characteristics of limb developmental processes.

### Mesodermal, but not ectodermal, cells show morphogenic properties for main budoid morphology

Next, we aimed to leverage budoids as a simplified system to dissect the effect of signaling center AER-like cells. For this, we first established a cell sorting strategy (Figure 4A, Figure S9), informed by the scRNA-Seq dataset (Figure S9A), to enrich mesodermal (EpCAM^Neg^), surface ectoderm-like (EpCAM^High^/CD9^High^), or AER-like (EpCAM^High^/CD9^Low^) cells. Enriched AER-like and mesoderm populations expressed the markers *Fgf8* and *Prrx1*, respectively (Figure S9D), and they were devoid of the pluripotent stem cell marker *Nanog*, while ectodermal-enriched populations displayed some expression of endoderm-associated *Sox17* (Figure S9E). These results established a successful enrichment strategy with minimal contaminant populations for cell type-specific assessments.

**Fig 4.**
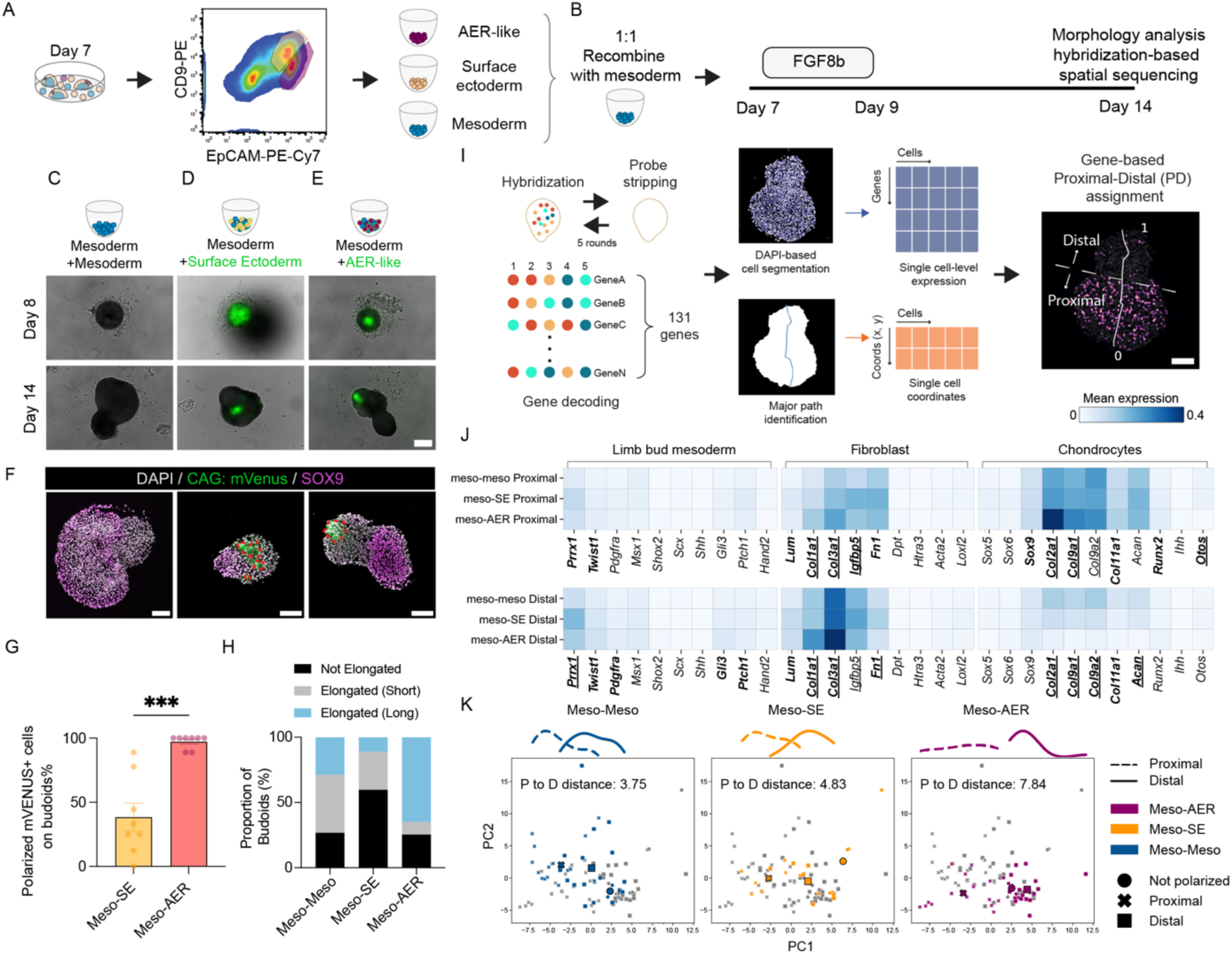
Recombinant budoids and *in situ* gene expression profiling reveal the impact of AER-like cells on cell identity and spatial organization. (A) Schematic showing an example of a fluorescence-activated cell sorting (FACS) strategy to enrich specific cell types. (B) The experimental design for recombinant budoids is outlined and followed by assays for morphological changes and spatial sequencing. (C-E) Overlaid brightfield and fluorescent images taken on days 8 and 14 show (C) mesoderm recombined with mesoderm, (D) mesoderm recombined with surface ectoderm-like, and (E) mesoderm recombined with AER-like cells. Please note the surface ectoderm-like and AER-like cells are derived from the *CAG:mVenus* line, serving as a fluorescent marker for tracking. Non-aggregated and shed cells, especially in ectodermal cells, are seen on day 8 samples. Distinct polarized *mVenus* positive AER-like cells are seen on day 14 in E. Scale bar = 100 μm. (F) Representative confocal images of sectioned recombinant budoids. mVENUS+ ectodermal cells are circled with a red dashed line. Please note distinct polarized mVENUS+ AER-like cells. Magenta: SOX9; Green: *CAG:mVenus*: Hoechst: DAPI. Scale bar = 250 μm. (G) A bar plot showing polarized *mVenus* group of cells on recombinant budoids. Each dot represents the average phenotype per replicate. Total number of budoids analyzed for Meso-SE n= 60, N=8; Meso-AER n=63, N=8. *** denotes *p*<0.001. (H) A bar chart with a weighted average of the proportion of recombinant budoids for their elongation. Total number of budoids analyzed for Meso-Meso n= 80; Meso-SE n=78, Meso-AER n= 63, from N=8. (I) Schematics describing the hybridization-based in situ sequencing (HybISS) methodology and subsequent analysis, leading to the unbiased identification of proximal and distal domains of recombinant budoids. Please see methods for details. The major axis is labeled with a line. From 1 to 0 represents distal to proximal directions. The dashed line splits the recombinants into distal and proximal domains. Scale bar = 100 μm. (J) The heatmap displays average expression levels of genes associated with limb bud mesoderm, fibroblasts, and chondrocytes in the recombinant budoid single cells within proximal and distal domains. Genes with statistically significant differences are highlighted, with bold indicating a significant difference between mesoderm-AER and mesoderm-mesoderm comparisons, while underlining signifies significant differences between mesoderm-AER and mesoderm-surface ectoderm. The number of HybISS sections analyzed for Meso-Meso n=20, N=4, Meso-SE n=12, N=3, Meso-AER n=20, N= 4. (K) Principal component analysis (PCA) of sectioned recombinant budoids illustrates the separation between the proximal and the distal domains. Proximal domains of individual recombinant sections are denoted by **X**, and distal domains are denoted by **■**. Sections without polarization are denoted by **●**. Density plots showing the distribution of samples with respect to PC1 are shown at the top of the PCA plots. The distributions of distal and proximal samples are represented with a straight line and a dashed line, respectively. The distance between the centroids of the distal and proximal samples (black-framed bold symbols) was quantified and displayed. Samples are color-coded. The number of HybISS sections analyzed for Meso-Meso n=20, N=4, Meso-SE n=12, N=3, Meso-AER n=20, N= 4.

We then tested the generation of budoids from only mesoderm, AER-like, or surface ectoderm-like cells and found that they all had disintegration and aggregation problems (Figure S10A). However, generating aggregates with a brief FGF8b exposure revealed that mesoderm-enriched cells alone can break symmetry and elongate (Figure 4C, Figure S10B-C), although they showed variability in size across experimental sets (Figure S10B). In contrast, the AER-like enriched population did not properly aggregate (Figure S10A-B), albeit there was a minor improvement with brief FGF8b treatment (Figure S10C). These phenotypes aligned with AER-like cells tested by sorting and aggregating day 7 *Fgf8:tdTomato*+ cells (Figure S9F, 10D). Surface ectoderm-like enriched cells were able to aggregate more with a brief FGF8b treatment but did not show symmetry breaking or elongation (Figure S10C). In summary, we concluded that mesoderm-enriched cells can display morphogenesis by breaking symmetry and elongation upon brief exposure to FGF8b, whereas ectodermal cells alone do not have a similar morphogenic capacity.

### Recombinant budoids reveal the impact of signaling center AER cells on budoid morphology and spatial organization

Then, we devised “recombinant budoids” to uncover the impact of signaling center AER-like cells on the mesoderm-enriched population (Figure 4B). Recombinants were generated by combining equal numbers of FACS-sorted mesodermal cells and AER-like cells. Similarly, we recombined mesoderm with surface ectoderm-like, or mesodermal cells as controls. Moreover, we generated AER-like and surface ectoderm-like cells from a constitutively active *mVenus*-expressing line to trace them. Due to aggregation problems (Figure S10A-E), all conditions received FGF8b treatment for 2 days and were grown for an additional 5 days before being examined for morphological features (Figure 4B).

Although we seeded the same number of cells for recombination, we observed different-sized aggregates at 1 day post-seeding in all conditions (Figure 4C-E, Figure S11A), possibly because of cell type-specific aggregation capacities (Figure S10A-B, Figure S11A). Mesoderm with AER-like recombination had a smaller area and contained fewer mVENUS+ cells at day 1 compared to other recombinants (Figure 4C-E, Figure S10F, and Figure S11A). Recombined AER-like or surface ectoderm-like cells did not expand and were further eliminated from the structures over time (Figure 4D-E, Figure S10F), consistent with the reduction of ectodermal cells in previous budoids experiments (Figure S7B, Figure S8C-D). Moreover, these ectodermal cells were typically seen together and engulfed in the budoids (Figure 4F), probably due to cell sorting and differential adhesion properties favoring ectoderm internalization, as suggested before^29^. Remarkably, 97.2(±1.8)% of the analyzed recombinant budoids had polarized mVENUS+ AER-like cells, while the same phenotype was only seen in 38.4(±10.9)% of mesoderm with surface ectoderm-like recombinations (Figure 4D-E, G, and Figure S10F). We then evaluated the fold change in area from initial aggregates to final structures as a proxy for their growth. All conditions showed comparable values (Figure S11A), indicating that AER-like recombination in this setup does not significantly impact the growth rate.

Importantly, mesoderm with AER-like recombinant budoids had substantial symmetry breaking and elongation compared to mesoderm with surface ectoderm recombinants (Figure 4H, Figure S11B), which remained in a more circular morphology (Figure 4D, Figure S10F). Given our results suggesting chondrogenesis is associated with symmetry breaking (Figure 3I, Figure S7B), the inability of surface ectodermal-like recombination to break symmetry and elongate may be consistent with the previously proposed function of ectodermal cells in counteracting chondrogenesis^4,30^. However, we did not observe a similar phenotype with the AER-like recombination (Figure 4H, Figure S11B). To determine whether this was due to the polarized spatial segregation of AER-like cells (Figure 4E, Figure S10F) and their potential local impact, we evaluated the chondrogenesis marker SOX9^31^, and the fibroblast/limb bud mesoderm marker TWIST1^32^. These markers mostly showed a mutually exclusive expression pattern across all samples, constituting most of the cells in structures (Figure S11C). Abundant SOX9+ cells were seen in mesoderm-enriched samples (Figure 4F, Figure S11C). Strikingly, AER-like cells and SOX9+ cells were mostly polarized at opposite ends of the structures (Figure 4F, Figure S11C), suggesting local suppression while permitting this differentiation in a distance. In contrast, mVENUS+ surface ectoderm-like cells could be found with some SOX9+ cells in close proximity without clear polarization (Figure 4F, Figure S11C). Thus, these recombination experiments confirmed the functional properties of ectodermal cells, consistent with previous suggestions that ectoderm impairs chondrogenesis^4,30^. Moreover, our experiments also distinguished specific effects of AER from surface ectoderm: AER-like cells impaired chondrogenesis only for neighboring cells without disrupting mesodermal morphogenetic capacity, whereas surface ectoderm recombination directly reduced chondrogenesis-based symmetry breaking.

Overall, our recombinant budoid strategy allowed us to reveal cell-cell interactions that are difficult to dissect *in vivo,* and how the ectoderm influences the spatial segregation of cell fate decisions and tissue organization.

### AER-like cells in recombinant budoids promote nearby mesoderm and fibroblast identities and establish tissue polarization permitting distant cartilage formation

We then investigated the molecular characteristics of our recombined structures using quantitative hybridization-based *in situ* sequencing (HybISS)^33^ assaying 131 limb development-related cell type and signaling pathway genes (Figure 4I, Figure S12). Leveraging the single-cell resolution of HybISS alongside spatial cellular coordinates, we separated individual samples into two domains (Figure 4I). Across all recombinants, we defined one side of the structures showing enrichment for chondrogenic markers (e.g., *Sox9, Col2a1, Col9a1, Col9a2*) (Figure 4K, Figure S13A), as the proximal domain. Meanwhile, the opposite side, the distal domain, showed higher expression of limb bud mesoderm, fibroblast (e.g., *Prrx1, Msx1, Twist1, Lum, Col1a1, Col3a1*) (Figure 4J, Figure S13A), and signaling genes (e.g., *Bmp4, Bmp7 – Id2, Id3*; *Wnt5a, Wnt6, Rspo2, Rspo3 -Axin2, Tcf7*; *Fgf7 -Dusp1, Spry2, Spry4*) (Figure S13B-F). Therefore, recombinant budoids have two distinct domains that resemble spatial features of limb buds: the proximal domain enriched for cartilage identity, and the distal domain with limb bud mesoderm and fibroblast states and high signaling activity.

Focusing on ectodermal cells, we found a few distally enriched potential remnants of AER-like cells (*Krt5*+), while surface ectodermal-like (*Krt8+/18+*) cells showed no clear enrichment for a specific location (Figure S14A). Previously reported AER-induced genes such as *Msx1, Bmp4, Wnt5a, Gli3,* and *Axin2*^27^ were also enriched at the distal domains of AER-recombinants (Figure S14B), further confirming the functionality of generated cells and their lasting effect.

AER-like with mesoderm recombinants showed high expression for fibroblast (*Prrx1, Twist1, Lum, Col3a1, Igfbp5, Col1a1*), and limb bud mesoderm genes (*Fgf10*, *Msx1*, and *Gli3*) across both domains with distal enrichment (Figure 4J, Figure S13A). In contrast, these signatures were either reduced or at comparable levels in surface ectoderm with mesoderm recombinants and even more significantly reduced in mesoderm-only recombinants (Figure 4J, Figure S13A). Instead, mesoderm-only recombinants had high chondrogenesis gene expression (e.g., *Col2a1, Acan, Col9a1, Col9a2*) in both domains (Figure 4J, Figure S13A), further supporting the notion that AER-like cells in budoids influence the patterned expression of chondrogenesis genes.

Finally, performing principal component analysis on samples classified into proximal and distal domains identified that the AER-like cells increased polarization between domains, while the other conditions showed more mixing (Figure 4K). Altogether, these results uncovered that signaling center AER-like cells promote or sustain the nearby limb bud mesoderm and fibroblast identities, while suppressing nearby chondrogenesis, and affect spatially distant cartilage formation by promoting polarization.

## Discussion

Here, we present a highly scalable and novel deconstructed approach to study signaling centers and limb morphogenesis. The budoid methodology provides a unique window into the complex interplay between the specialized signaling center AER and other limb development-related cell populations, amenable to capturing quantitative cell-cell interactions, which are challenging to study *in vivo*. Taking advantage of this platform, we have uncovered that mesodermal cells possess the capacity to self-organize and break symmetry upon brief FGF8b treatment, independently of any 3D embedding (e.g., Matrigel), and that even a transient and small number of AER-like cells can exert morphological, spatial, and molecular changes on mesodermal cell fates. Critically, these findings also show that the impact of specialized signaling centers is not only limited to nearby cells but can extend to spatially distant populations, consequently shaping tissue organization.

Although our initial objective was to generate only signaling center AER cells, our protocol unexpectedly produced self-organizing heterogeneous cultures with limb development properties. It remains unclear how our protocol led to the simultaneous formation of these populations. Nonetheless, this strategy allowed us to systematically study individual limb populations beyond just signaling center AER cells, and to develop a new simplified model showing aspects of limb morphogenesis. In contrast to earlier limb models (including a limb bud organoid-like model)^9–13^, we characterized our system by comprehensive single-cell analyses and comparisons with *in vivo* samples, examined the molecular and cellular properties of each population separately, and performed systematic recombination experiments with quantitative spatial gene expression profiling to characterize their functionality. Importantly, the generation of budoids is highly feasible, inexpensive, and efficient, with ∼80% of aggregates exhibiting chondrogenesis-based symmetry breaking, suitable for exhaustive biological inquiry. Altogether, these advances position budoids as a multipurpose and easily adoptable method for a wide range of studies, accessible to many laboratories.

Our stem cell-based protocols can lead to mechanistic and quantitative understanding of specialized signaling centers and limb morphogenesis, that can also guide regenerative therapies. Bioengineering strategies have been employed to emulate single or multiple morphogen gradients^34,35^, and our generated *in vivo*-like specialized signaling center cells could enable the dissection of the matrix of signaling gradients and cell-cell interactions, potentially also improving endeavors of computational limb modeling^36^. Furthermore, our protocols can be adapted to stem cells from other species, including human, for comparative analyses ^37^. Lastly, the scalability of budoids paves the way for high-throughput genetic and chemical screens to identify factors that promote cartilage formation for therapeutic purposes.

Efforts to regenerate mammalian limbs or digits have focused on transplanting limb bud mesoderm with limited success^9,38^. Meanwhile, the effectiveness of specialized signaling centers^39^, and particularly AER cells^40,41^, has been emphasized in amphibian regeneration. Therefore, our methods robustly forming specialized signaling center AER-like cells, and other limb cell types could be foundational in fundamental regeneration research, as well as transplantation studies aimed at inducing blastema for mammalian regeneration. On this basis, several important areas for improvement remain. Firstly, our protocol produces newly forming AER-like cells, awaiting identification of factors to maturate them. This can also inform how AER is regulated *in vivo*, as well as examine their possible formation following limb amputations. Secondly, our functional assays demonstrate that AER-like cells are hard to maintain and show significant aggregation problems, which effectively limits transplantation studies. Finally, although we were able to FACS-enrich specific populations, refinements to our protocol will increase their purity. Due to the highly efficient nature of our method, we anticipate that future studies can systematically address these challenges.

In summary, we provide a new methodology for studying specialized signaling centers and limb morphogenesis in a simplified setting. Our methodology could lead to an in-depth and quantitative understanding of development, congenital disorders, cross-species features, and new directions for regenerative medicine.

## Acknowledgments

We thank Fides Zenk and Alfonso Martinez-Arias for discussions; Life Science Editors for editing services; the EPFL Gene Expression Core Facility for supporting this work on 10X-Genomics and sequencing library preparations; the EPFL BioImaging and Optics Facility and the EPFL Flow Cytometry Facility for their technical assistance; the EPFL Protein Production and Structure Core Facility for providing LiF; and the EPFL Center of PhenoGenomics for mouse housing and uterus dissections. C.A. is supported by EPFL School of Life Sciences ELISIR Scholarship, the Foundation Gabriella Giorgi-Cavaglieri, Branco Weiss Fellowship, SNSF NRP79 (407940-206349), and Novartis Foundation for Medical-Biological Research.

## Author contributions

Conceptualization: Mainly C.A., with some help from E.S. and J.Z.;

Experimentation: Mainly E.S., with C.A., O.K., K.H., G.T., L.C., and A.I. contributing to various experiments, G.T. and K.H. performed HybISS experiments, T.H. helped with the initial HybISS experiment;

Computational Analysis: Mainly J.Z., K.H. performed the morphology analysis, A.D. helped with the initial HybISS analysis, H.J. helped with the initial morphology analysis;

Data interpretation: Mainly C.A., with E.S. and J.Z. also helping;

Funding acquisition: C.A.;

Project administration: C.A.;

Supervision: C.A.;

Writing -initial draft: Mainly C.A. with help from E.S. and J.Z;

Writing -review and editing: All authors;

## Data and materials availability

Sequencing data are available on GEO database under accession number XXX (available to reviewers upon request and will be made public upon publication). Processed scRNA-seq and HybISS data can be downloaded from Zenodo (https://zenodo.org/records/10961079). Analysis scripts are available at https://github.com/AztekinLab/Budoids_2024. Requests for materials should be directed to C.A.

## Methods and Protocols

### Mouse embryonic stem cell culture

129/SvEv mouse embryonic stem cells (mESCs) (a kind gift of Denis Duboule) were maintained in 6-well plates coated with 0.1% gelatin (Merck, ES-006-B, and Stem Cell Technologies, 07903 used interchangeably) in serum + 2i/LiF (DMEM, high glucose, Glutamax supplement (Gibco, 61965-026) supplemented with 10% ES cell FBS (Gibco 16141-079), 1X non-essential amino acids (Gibco, 11140035), 1 mM sodium pyruvate (Gibco, 11360-039 or 11360070), 0.1 mM 2-mercaptoethanol (Gibco, 31350-010), 100 U/mL penicillin/streptomycin (Gibco, 15140122), 100 ng/mL LiF (produced by the Protein Production and Structure Core Facility at EPFL), 1 μM PD0325901(Sellechckem, S1036), (3 μM CHIR99021 (ApexBio (A3011) or Calbiochem (361559) used interchanably). Media were changed every 2 days, or cells were passaged every 2-4 days. Cells were grown in an incubator at 37°C and 5% CO2. Routine mycoplasma tests were performed.

### Plasmid construction and cell line establishment

A CRISPR-Cas9 approach was used to generate the *Fgf8* reporter line. Annealed oligos encoding gRNAs targeting near the *Fgf8* stop codon (the forward oligo 5’-caccGAGCGCCTATCGGGGCTCCG-3’, the reverse oligo 5’-aaacCGGAGCCCCGATAGGCGCTC-3’) were ordered from IDT and ligated into pX330 vector digested with BbsI as described previously^42^. The 5’ homology arm (800 bp) and 3’ homology arm (800 bp) flanking from the *Fgf8* stop codon were amplified by PCR from mouse genomic DNA and cloned into the homology-directed repair donor vector containing tdTomato and puromycin resistance gene (a kind gift of Naoko Irie)^43^ using in-fusion cloning (Clontech) according to the manufacturer’s recommendations. The cloned plasmid was verified by Sanger sequencing. For knock-in, 20,000 mESCs were seeded in 24-well plates coated with 0.1% gelatin. One day after seeding, mESCs were transfected with donor vector and gRNA-containing pX330 vector using 2.4 μL Lipofectamine 2,000 (Invitrogen ref: 11668-019) per 800 ng total plasmids according to the manufacturer’s recommendations. Puromycin selection was applied. The remaining cells were passaged, and 1,000 cells were seeded on 60mm x 15mm plates coated with 0.1% gelatin for colony picking. Grown single colonies were picked manually, expanded, and genotyped by PCR. Genomic DNA was first extracted from cell pellets in lysis buffer (10 mM Tris pH 8.0, 100 mM NaCl, 10 mM EDTA, and 0.5% SDS; proteinase K (PK) was added immediately before lysis at a final concentration of 0.2 mg/ml) at 56 °C for 4 hours to overnight, followed by PK inactivation at 98 °C for 10 min ^43^. The supernatant was used directly in genotyping PCR. PCR-verified clones were expanded, and genomic DNA from the expanded line was used to confirm integration by Sanger sequencing. Primers used for genotyping are shown in Supplementary Table 4. No leaky expression of *TdTomato* was observed in mESCs.

PiggyBac (PB) transposase-based integration was used to generate *CAG:mVenus* and *Prrx1^Enh^:mVenus* mES cell lines. The PB vector containing *CAG:mVenus* was kindly provided by Naoko Irie. The *Prrx1^enh^* ^24^ is cloned from mouse genomic DNA into the PB vector using In-Fusion (Takara) according to the manufacturer’s recommendations. The cloned plasmid was verified by sequencing. For integration, 20,000 mESC cells were seeded in gelatinized 24-well plates. One day later, cells were co-transfected with one of the PB vectors, puromycin resistance plasmid (a kind gift of David Suter), and PiggyBac transposase plasmids using 2.4 μL Lipofectamine 2000 (Invitrogen ref: 11668-019) per 800 ng total plasmids according to the manufacturer’s recommendations. Puromycin selection was performed. No leaky expression of *Prrx1^Enh^:mVenus* was observed in mESCs.

### Heterogenous culture induction

The mESCs were washed with 1X PBS (Gibco, 10010015), dissociated with 1X 0.25% trypsin-EDTA (Gibco, 25200072 for 2 minutes at 37°C, and de-activated in mESC media without inhibitors (2i/Lif). Cells were collected in 15 ml falcons containing 8 ml 1X PBS (to dilute the remaining factors further) and centrifuged at 137 rcf for 5 minutes at RT. The media was aspirated and cells were resuspended to single cells in non-neural ectoderm, NNE, media (GMEM (Gibco, 11710-035) supplemented with 1.5% knockout serum replacement (Gibco, 10828010), 1X non-essential amino acids (Gibco, 11140050), 1 mM sodium pyruvate (Gibco, 11360-039 or 11360070), 0.1 mM 2-mercaptoethanol (Gibco, 31350-010), 0.1 mg/mL Normocin (InvivoGen, ant-nr-1) final concentrations) and counted. 40,000 cells were seeded in 12-well plates coated with 0.1% gelatin and incubated at 37°C in 5% CO2. On day 1 after seeding, the cells were washed with 1X PBS, and the medium was changed to fresh NNE medium (1 mL per well). On day 3, significant cell death was observed. The medium was aspirated, and the cells were washed twice with 1X PBS and changed to fresh 1 ml NNE containing SB431542 (Apexbio, A8249) at 1 μM concentration and 100 or 200 ng/mL BMP4 (PeproTech, 120-05ET) final concentrations. On day 5, the cells were washed twice with 1X PBS, and the medium was changed to fresh 1 ml NNE containing 100 ng/mL FGF10 (PeproTech, 100-26), 100 or 200 ng/mL BMP4 (PeproTech, 120-05ET) and 4.5 μM CHIR99021 (Apex Bio) or 3 uM CHIR99021 (Calbiochem) final concentrations. Media changes were performed at 48-hour intervals (±2 hours). All recombinant proteins and chemicals used in cell culture were reconstituted in the buffers as recommended by the supplier. During this study, we detected batch variation for CHIR99021 and recombinant BMP4. The majority of the experiments in this study were performed with 4.5 μM CHIR99021 (Apex Bio), and all experiments (except 2D culture scRNA-seq experiments) were replicated with 4.5 μM CHIR99021 (Apex Bio). Similarly, the majority of the experiments in this study were performed with 100 ng/mL BMP4 (PeproTech, 120-05ET), and all experiments were replicated with 100 ng/mL BMP4 (PeproTech, 120-05ET).

### Quantitative RT-PCR

For quantitative RT-PCR (qRT-PCR) analysis, total RNA was purified using the RNeasy Mini Kit (Qiagen, 74106) for day 7 samples, and PicoPure RNA isolation kit (Thermo, KIT0204) for sorted samples according to the manufacturer’s protocol for both kits. 1 μg of RNA was used for cDNA synthesis using SuperScript II (Invitrogen, 18064-014). qRT-PCR was performed with Power SYBR Green PCR Master Mix (Applied Biosystems, 4367659) using the QuantStudio 6 or 7 Flex Real-Time PCR Systems (Applied Biosystems), and data were calculated using the delta-delta Ct method. Rpl27 was used as a housekeeping gene in all experiments. In Figure S1E, L, the relative expression of each gene at day 7 was calculated relative to the mESCs (day 0) used in the same experiment. In Figure S9D, E, the relative expression of sorted populations was calculated relative to the mRNA extracted from one of the mESCs (day 0) data. Hamilton Microlab Star (Hamilton) was used to assemble the plates. Only two technical replicates were used for each sample, as the Hamilton Microlab allows precise pipetting. The primers used in qRT-PCR are shown in Supplementary Table 4.

### Generation of budoids in 3D cultures from mouse embryonic stem cells

To generate budoids, the first 7 days followed the induction protocol as described above. On day 7, cells were dissociated with 0.25% trypsin-EDTA (Gibco, 25200072) for 4-5 minutes at 37°C, and deactivated with mESC media without inhibitors (2i/Lif). Cells were collected in a 15 ml falcon containing 8 ml 1XPBS and centrifuged at 400 rcf for 5 minutes, resuspended in NNE medium, and counted. 90,000 cells/ml suspension was prepared with NNE medium containing 300 ng/mL WNT3a (PeproTech, 315-20) and 450 ng/mL FGF8b (PeproTech, 100-26), and 100 μL of cell suspension per well was seeded into 96-well low attachment plates (Thermo Scientific, 174929). Please note that in the experiments for Figure S5, different concentrations of these recombinant proteins were tested. For the BMP pathway perturbation experiments described in Figure 3I and J, LDN193189 (MedchemExpress, HY-12071A-10MM/1ML), recombinant NOGGIN (PeproTech, 120-10C), or recombinant BMP4 (PeproTech, 120-05ET) were added at specified concentrations during budoid generation. In these experiments, the final concentration of LDN193189, NOGGIN, or BMP4 was maintained throughout the experiments. 0.4% DMSO was used as a vehicle control. No reservoirs were used to seed the cells to eliminate the possibility of WNT3a adhering to the reservoir surface. 50 μL of the media was changed once every 48 hours (±2 hours), and samples were collected at the reported time points. Cells were grown in an incubator at 37°C and 5% CO2. The outer layer of the 96-well plates was not used during budoid generation and was filled with 100 μl of 1XPBS to avoid side effects due to evaporation.

### Generation of budoids from *in vivo* E12.5 mouse limbs

Mice embryo samples were collected in accordance with the Swiss Federal Veterinary Office guidelines and as authorized by the Cantonal Veterinary Office (cantonal animal license no: VD3652c, and national animal license no: 33237).

Pregnant female CD1 mice were purchased from Charles Rivers Laboratories, and E12.5 embryos were collected by dissecting the uterine horn. The hindlimbs and forelimbs of embryos were collected and dissected for proximal and distal regions, as shown in Figure S6A. No distinction was made between left or right hind or forelimbs, and no sex determination was done. Samples were collected in tubes containing 1XPBS, each containing 10-15 samples. Samples were washed once with 1X PBS, and dissociated with 0.25% trypsin-EDTA (Gibco, 25200072) for 5 minutes at 37°C, deactivated with DMEM (with 10% FBS). Cells were collected and centrifuged at 400 rcf for 5 minutes and resuspended in NNE medium containing no additives, 450 ng/mL FGF8b (PeproTech, 100-26), or 300 ng/mL WNT3a (PeproTech, 315-20) and 450 ng/mL FGF8b (PeproTech, 100-26). 100 μL of cell suspension per well was seeded into 96-well low attachment plates (Thermo Scientific, 174929). 50 μL of the media was changed once every 48 hours (±2 hours), and samples were collected at the reported time points. Cells were grown in an incubator at 37°C and 5% CO2. All recombinant proteins and chemicals used in the cell culture were reconstituted in the buffer as the supplier recommended.

### MOrgAna (Machine-learning Organoid Analysis)-based area and elongation analysis, and mVENUS-based polarization quantification

MOrgAna (Machine-learning Organoid Analysis)^44^ was used to automate the area and elongation phenotypes of the generated 3D structures. 5% of the generated data were used to train MORgAna by manual masking for each different experimental design. MOrgAna outputs Excel files with the area and major and minor axis lengths. For elongation, a set of structures was manually examined and classified into three categories: not elongated, elongated (short), and elongated (long) by two independent researchers. These classifications were compared to the major and minor axis measurements obtained from MOrgAna. The ratio of minor axis length to major axis length was taken and thresholds were determined that resulted in minimal classification errors for each category. Values ≥0.85 correspond to no elongation, 0.75-0.85 to (short) elongation, ≤0.75 to (long) elongation. To quantify polarized mVenus-positive cells in Fig 4G, brightfield overlaid with fluorescent images of recombinant budoids having clear mVenus signal was used, and the number of samples showing polarization was manually counted.

Enriched cells were counted and processed as described in the regular budoids generation protocol (with 90,000 cells/ml), with or without 450 ng/ml FGF8b treatment, indicated in the relevant figures. For recombinant budoids experiments, individual enriched populations were mixed in separate tubes at a 1:1 ratio, totaling 90,000 cells/ml with or without the addition of 450 ng/ml FGF8b, indicated in the relevant figures. 100 μl of the solution was seeded into 96-well low attachment plates (Thermo Scientific, 174929). Images were taken 1 (day 8 of the full protocol) and 7 (day 14 of the full protocol) days after seeding via Nikon Eclipse Ti2.

### Fluorescence-activated cell sorting (FACS), and flow cytometry-based quantification

For antibody labelling-based ectodermal cell sorting experiments, CAG:mVENUS mESCs line was used to sort surface ectoderm-like (CD9^High^/EPCAM ^High^) or AER-like (CD9 ^Low^/EPCAM ^High^) cells. Day 7 induction cells were washed once with 1X PBS, and dissociated with 0.25% trypsin-EDTA (Gibco, 25200072) for 5 minutes at 37°C, deactivated with DMEM (with 10% FBS). Cells were collected, centrifuged at 400 rcf for 5 minutes, and resuspended in cold FACS buffer (1X PBS supplemented with 1% BSA (Gibco, 15260037). Cells were then stained for the surface markers EpCAM (Invitrogen, 25-5791-80) and CD9 (Invitrogen, 12-0091-81) in FACS buffer for 45 minutes on ice in the dark. The cells were then washed with 1 mL of ice-cold FACS buffer and centrifuged at 600 rcf for 5 minutes, with the washes repeated a total of 2 times. Cells were then resuspended to a single cell suspension in a cold FACS buffer in a round bottom polystyrene tube, with cell strainer (Corning 352235), and stored in the dark until analysis/cell sorting. For EpCAM staining-based mesoderm enrichment (EpCAM^Neg^), wild-type mESCs were used with the above protocol, except CD9 staining was not performed during labeling. For *Fgf8:tdTomato*-based cell sorting, the same induction and dissociation protocols were repeated with *Fgf8:tdTomato* mESC line; Dissociated cells were resuspended to a single cell suspension in cold FACS buffer in a round bottom polystyrene tube, with cell strainer, and stored in the dark until analysis/cell sorting. FACSAria Fusion and Sony SH800 were used for cell sorting. Figure S9B, C, and F present example gating strategies for cell sorting. FACS experiments were also used to quantify EpCAM, and *Fgf8:tdTomato*-positive cell numbers, reported in Figure 1E-F. Figure S1J and K present examples of flow cytometry-based quantifications and gating. In all experiments, an unstained wild-type control was used to label negative cells, and single staining was performed for compensation when necessary. FlowJo was used for further analysis.

### Individual lineage and recombinant budoids experiments

The above-described cell sorting experiments were performed to enrich individual lineages. Enriched cells were counted and processed as described in the regular budoids generation protocol (with 90,000 cells/ml), with or without 450 ng/ml FGF8b treatment, indicated in the relevant figures. For recombinant budoids experiments, individual enriched populations were mixed in separate tubes at a 1:1 ratio, totaling 90,000 cells/ml with or without the addition of 450 ng/ml FGF8b, indicated in the relevant figures. 100 μl of the solution was seeded into 96-well low attachment plates (Thermo Scientific, 174929). Images were taken 1 (day 8 of the full protocol) and 7 (day 14 of the full protocol) days after seeding via Nikon Eclipse Ti2.

### Immunostaining and hybridization-chain-reaction (HCR)

For immunostaining in 2D cultures, the induction protocol was performed in 12-well glass-like polymer coverslip bottom plates (Cellvis P12-1.5P). On day 7, samples were washed with 1X PBS and fixed with 4% formaldehyde (FA, Thermo Fisher Scientific,119690010) for 10-12 minutes at RT. This was followed by washing samples with 1X PBS for 2 x 5 minutes. To permeabilize cells, samples were incubated with 0.1% Triton X-100 (Sigma-Aldrich) in 1X PBS (hereafter, PBS-T) for 3 ×10 minutes. The samples were then incubated with blocking solution (50% Cas-Block (Thermo, 008120) and 50% PBS-T) for 30 minutes. The samples were then incubated with primary antibodies (TP63 (Abcam, ab735), TFAP2C (Cell Signaling, 2320S), LEF1 (Abcam, ab137872), diluted in blocking buffer overnight at 4°C. To enhance the reporter signal, suitable samples were stained with anti-GFP (Abcam, ab13970) (for Prrx1 enhancer reporter experiments) or anti-RFP (a kind gift from Sebastian Pons ^45^) (for Fgf8 reporter experiments). The next day, samples were incubated with 1X PBS-T for 3×10 minutes. Samples were then blocked by blocking solution for 30 minutes, followed by incubation with blocking solution containing Alexa Fluor (AF-488 (Thermo, A11039, A21202), 594 (Thermo, A11012) and/or 647 (Thermo, A32728)-conjugated secondary antibodies specific for the host species of the primary antibodies for 1 hour at RT in the dark. During secondary antibody incubations, Phalloidin staining was done by adding the relevant material (Abcam, ab176757, ab176753) for relevant experiments. After secondary antibody incubations, samples were washed with PBS-T for 3 x 10 minutes and then with 1X PBS for 3 x 10 minutes. Samples were incubated in 1X PBS with Hoechst 33342 (Sigma, 2261) for 15 min to stain for nuclei and then were washed with 1X PBS. The protocol was performed at room temperature (RT) unless otherwise stated. The samples were stored at 4°C in the dark (up to 2 weeks) before imaging on a Leica SP8 inverted confocal microscope. All incubations were performed on a gentle rotation system.

For whole mount budoids immunostaining, budoids were collected from 96-well plates using low binding tips (Sigma-Aldrich, Z719668), with a cut tip, and handled with low binding tips throughout the protocol. Collected budoids were washed with 1x PBS, and fixed with 4% formaldehyde (FA, Thermo Fisher Scientific,119690010) in 1X PBS at room temperature. This was followed by incubating budoids with 1X PBS for 10 minutes. To permeabilize samples, budoids were incubated with PBS-T for 3 x 10 minutes. Budoids were then incubated in a blocking solution (50% Cas-Block (Thermo, 008120) and 50% PBS-T) for 30 minutes. Budoids were then incubated with primary antibodies (TP63 (Abcam, ab735), SOX9 (Merck, AB5535), E-CADHERIN (BD, 610405)) diluted in a blocking buffer overnight at 4°C. The next day, budoids were incubated in PBS-T for 3×10 minutes, re-blocked with blocking solution for 30 minutes, and incubated with blocking solution containing Alexa Fluor (AF-488 (Thermo A32731), and/or 647 (Thermo, A32728)-conjugated secondary antibodies specific for the host species of the primary antibodies for 1 hour at RT in the dark. After secondary antibody incubations, budoids were washed with PBS-T for 3×10 minutes and then with 1X PBS for 3×10 minutes. Budoids were incubated in 1X PBS with Hoechst 33342 (Sigma, 2261) for 15 min at RT to stain for nuclei and then were washed with 1X PBS The samples were stored at 4°C in the dark (up to 2 weeks) before imaging on a Leica SP8 inverted confocal microscope. Please note that for E-cadherin or TP63 staining, samples were fixed with 4% formaldehyde 20 min and overnight, respectively, and the same above protocol was applied. All incubations were performed on a gentle rotation system to prevent potential buckling of budoids. The protocol was performed at room temperature (RT) unless otherwise stated.

Hybridization chain reaction (HCR)^46^ was performed as described previously for 2D cultures, and with modifications for budoids, and materials for HCR, including probes, were purchased from Molecular Instruments. Budoids were collected from 96-well plates using low binding tips (Sigma-Aldrich, Z719668) with a cut tip and handled with low binding tips throughout the protocol. Budoids were washed with 1X PBS and then fixed with 4% formaldehyde (FA, Thermo Fisher Scientific, 119690010) in 1X PBS for 30-60 min. Samples were permeabilized in 70% ethanol for 1 hour and collected in Eppendorfs. The supernatant was removed, washing solution was added, and samples were rotated for 10 minutes. The supernatant was removed and replaced with hybridization buffer for 30 min incubation at 37°C. In parallel, probe solution was prepared by diluting mRNA targeting probes to 30-40 nM in 200 μl hybridization buffer and incubated at 37°C for 30 min. The hybridization buffer was removed from the samples, and the probe solution was applied to the samples for 12-16 h incubation at 37°C. The samples were then washed twice for 10 minutes with wash buffer and twice for 20 minutes with 5× SSC-T at room temperature. To visualize the probes, an amplification solution was prepared by first heating the pairs of fluorophore-tagged hairpins (h1 and h2 hairpins) corresponding to the probes to 95°C for 90 s. The hairpins were then left in the dark at room temperature for 30 min. The final amplification solution was prepared at 40-60 nM h1 and h2 in 200 μL amplification buffer. Samples were first incubated in an amplification buffer without hairpins for 10 min, then placed in the final amplification solution at room temperature, protected from light, for 12-16 h. Samples were washed for 2×20 min with 5× SSC-T. The samples were stained with 1X PBS with Hoechst 33342 (Sigma, 2261) and washed in 1X PBS for 10 min. Stained budoids were transferred to imaging plates and stored at 4°C in the dark (up to 2 weeks) before imaging on a Leica SP8 inverted confocal microscope. These procedures were performed at room temperature, unless otherwise stated.

In Figure S1F, 2D cultures were first processed with HCR and then with immunostaining as described above with minor modifications. Briefly, samples were processed with HCR (as reported before^46^). Then, the samples were incubated with blocking solution (50% Cas-Block (Thermo, 008120) and 50% PBS-T) for 30 minutes. The samples were then incubated with TP63 primary antibody (Abcam, ab735) diluted in blocking buffer overnight at 4°C. The next day, samples were incubated with 1X PBS-T for 3×10 minutes. Samples were then blocked by blocking solution for 30 minutes, followed by incubation with blocking solution containing Alexa Fluor (AF-680 (Thermo, A32729)-conjugated secondary antibodies for 1 hour at RT in the dark. After secondary antibody incubations, samples were washed with PBS-T for 3 x 10 minutes and then with 1X PBS for 3 x 10 minutes. Samples were incubated in 1X PBS with Hoechst 33342 (Sigma, 2261) for 15 min to stain for nuclei and then were washed with 1X PBS. The protocol was performed at room temperature (RT) unless otherwise stated. The samples were stored at 4°C in the dark (up to 2 weeks) before imaging on a Leica SP8 inverted confocal microscope.

For immunostaining recombinant budoids in Figure 4, and Figure S11, flash-frozen O.C.T. embedded samples were used. Please note that the subsequent sections of HyBISS samples were used in these experiments. The sections were allowed to acclimate to room temperature and fixed with 4% paraformaldehyde for 15min (Electron Microscopy Sciences, 15710-S). This was followed by one wash in 1XPBS, and the samples were incubated in PBS-T for 3 x 10 minutes. The samples were then incubated with a blocking solution for 1 hour at RT and they were then incubated with primary antibodies (SOX9 (Merck, AB5535), TWIST1 (Abcam, ab50887)) diluted in a blocking buffer overnight at 4°C. To enhance GFP signal, suitable samples were stained with anti-GFP (Abcam, ab13970). The next day, they were washed three times with the wash buffer (0.1% Triton X-100 in PBS) and incubated with blocking solution (50% Cas-Block (Thermo, 008120) and 50% PBS-T) for 15 minutes. Then the sections were incubated with Alexa Fluor (AF-488 (Thermo, A11039), 594 (Thermo, A11012) and/or 647 (Thermo, A32728)-conjugated secondary antibodies specific for the host species of the primary antibodies for 1 hour at RT in the darkAfter three washes in wash buffer and then in 1XPBS sections were mounted with either SlowFade Gold Antifade Mountant with DAPI (Invitrogen S36939) or VECTASHIELD antifade mounting medium with DAPI (VectorLabs, H-1200-10). The samples were stored at 4°C in the dark (up to 2 weeks) before imaging on a Leica SP8 upright confocal microscope. Images were analyzed using Fiji software.

### Confocal and brightfield imaging

Brightfield images in Figure S1A-D, Figure 2D, Figure S5B, E, and Figure S6E were taken with Olympus CKX53SF. Brightfield images in Figure S6A were taken with Nikon Stereo Microscope SMZ745T KL 300 LED Light Source. Brightfield and fluorescence images in Figure 2G, Figure 4, Figure S10 were taken with Nikon Eclipse Ti2. Confocal images in Figure 1, Figure S4, Figure 3, and Figure S8 were taken with a Leica SP8 inverted confocal microscope with 10x/0.30 HC PL Fluotar or 20x/0.75 HC PL APO objectives. Confocal images in Figure 4F and Figure S11 were taken with a Leica SP8 upright confocal microscope with 20x/0.75 HC PL APO or 40x/1.25 HC PL APO objectives. For the confocal images, LAS X was used to set up tiled images, and a 10-20% overlap between tiles was used. All images were analyzed using Fiji software. Fiji was used to maximize the projection of the z-stacks and to adjust the contrast to emphasize biological relevance. When necessary, images were cropped, flipped, and/or rotated to highlight biological relevance.

### SOX9 distribution quantification in budoids

Maximum projections for z-stack images of organoids were manually masked in Fiji, and fluorescence intensity was calculated across the organoid major axis using the built-in intensity plot profile tool. The grey values and length of each organoid were normalized to values between 0 and 1. Averaged intensity profiles were generated by bucketing using 200 bins for the organoid length at increments of 0.005.

### Sample preparation for single-cell mRNA sequencing

The 2D induction and budoids generation were performed as described above, and samples were collected at designated time points. In the first replicate of 2D cultures, samples were collected during day 0, day 3, day 5, day 7 of the protocol, dissociated with 0.25% trypsin-EDTA (Gibco, 25200056) for 5 minutes at 37°C, deactivated with mESC media without inhibitors (2i/Lif). Cells were collected, dissociated and centrifuged at 400 rcf for 5 minutes, resuspended in 1X PBS supplemented with 1% BSA (Sigma, A9418-10G), and filtered through a strainer. Samples were multiplexed using CellPlex, according to the manufacturer’s recommendations, scRNA-seq libraries were generated using 10X Genomics (v3 chemistry) and sequenced in pools of two samples per lane on an Illumina HiSeq 4000, with the following parameters: 28 bp -read 1; 8 bp -i7 index; and 91 bp -read 2, as per standard 10X Genomics recommendations. In the second replicate of 2D cultures, only day 5 and day 7 samples were collected and processed in the same way, sequenced on Novaseq 6000 with parameters: 28 bp -read 1; 8 bp -i7 index; and 91 bp -read 2. For budoids sequencing experiments, the first experiment contained 60 pooled D12 elongated budoids, and processed with CellPlex as described above. The second experiment contained 60 pooled D9 budoids and 60 elongated D12 budoids. Briefly, individual budoids were collected from 96-well plates with a low attachment tip with a cut tip, and washed with 1X PBS, dissociated with 0.25% trypsin-EDTA (Gibco, 25200072) for 10 minutes at 37°C with shaking at 300 rpm, deactivated with mESCs media without inhibitors (2i/Lif). Cells were collected, centrifuged at 600 rcf for 5 minutes at 4°C, resuspended in 1X PBS supplemented with 0.04 % BSA (Sigma, A9418-10G), filtered and counted. Then, scRNA-seq libraries were generated and sequenced. These samples were processed and sequenced on Novaseq 6000 with parameters: 28 bp -read 1; 10 bp -i7 index; and 90 bp -read 2. Both replicates of 2D cultures, and the first replicate of budoids received 3 uM CHIR99021 (Calbiochem) on day 5 of the induction, and the second replicate of budoids received 4.5 uM CHIR99021 (ApexBio) on day 5 of the induction, and the results showing similarity across replicates are shown in Figure S7C.

### Analysis of single-cell mRNA sequencing

Each experiment was processed separately with CellRanger (v7.1.0) to obtain the expression matrix. Cells were filtered with dataset-specific thresholds based on mitochondrial percentage and the number of UMI and genes. (Supplementary Table 1). The expression matrix was normalized. Technical confounding factors, including cell cycle effect and sequencing depth, were regressed out with ScaleData function in Seurat (v4.3.1) ^47^.

Seurat RPCA-based integration was used to perform the co-clustering of cells from different experiments. In the integration of the 2D cells (day 0, 3, 5, and 7) from experiments 1 and 2 (Figure 1G, Figure S2), genes were ranked by the number of datasets they are deemed variable in (*SelectIntegrationFeatures*) and the top 3000 genes were used to integrate all the datasets (*FindIntegrationAnchors* and *IntegrateData*). The first 30 principal components (PCs) were chosen to build a K nearest neighbors (KNN) graph and clustering. Cell-type annotation was based on the expression of literature-supported cell-type markers. UMAP was performed for visualization. The default setting was used unless noted. Similarly, cells from day 9 and 12 from budoids sequencing experiments 1, 2 and 3 were integrated (Figure 3B, Figure S7). Data were normalized using SCTransform, where cell cycle effect and sequencing depth were regressed out. The final UMAP was generated using the first 40 PCs.

### Transcriptome-wide comparison of cluster similarity between *in vivo* and *in vitro* cells

ClusterFoldSimilarity ^48^ was used to calculate the cluster similarity between *in vivo* tissue and our *in vitro* system (Figure S3B, Figure S8A). ClusterFoldSimilarity scores cluster similarity by comparing the fold-change patterns in clusters that share a common set of genes. The top 3000 highly variable genes identified by *Seurat::FindVariableFeatures* were used to calculate the similarity. Limb bud mesoderm subtypes were annotated based on a published comprehensive atlas ^22^.

### Single-cell gene set enrichment analysis (scGSEA) of cell fate decision genes

The potential differentiation direction of cells was determined with the AddModuleScore function in Seurat (Figure 3F) using gene lists from published literature (Supplementary Table 1)^26^. Briefly, the average expression levels of each gene list were calculated and subtracted by the aggregated expression of control genes that were randomly selected with similar expression levels. To determine the significance between genes/modules at different time points, the Wilcoxon Rank Sum test was performed.

### Sample preparation for hybridization-based in situ sequencing (HybISS)

For in vivo limbs, pregnant female CD1 mice were purchased from Charles Rivers Laboratories, and E12.5 embryos were collected by dissecting the uterine horn. Embryos were then placed in cold 1X PBS, and the hindlimbs were isolated using fine forceps and scissors. Immediately after, the limb tissues were placed in optimal cutting temperature (OCT) and flash-frozen in a bath with isopentane and dry ice. Tissues were then stored at −80 °C until sectioning. For cryosectioning, tissues were cut at 10 μm using a cryostat (Leica CM1950) and stored at −80 °C until HybISS processing. For recombinant budoids, the same protocol was performed instead of hindlimb samples.

### Hybridization-based in situ sequencing (HybISS)

131 genes were selected for HybISS to reflect the cell identity and critical pathways based on the literature (Supplementary Table 2)^26^. Padlock probes (PLPs) were designed using the pipeline https://github.com/Moldia/multi_padlock_design with default parameters. After target sequences were obtained, five targets were selected randomly per gene. If less than five targets were found, all the targets were selected. The backbone of the PLPs includes a 20 nucleotide ID sequence that is specific to each gene and a 20 nucleotide ‘anchor’ that is the same for all PLPs and served only as linker sequences in this study. The designed probes were ordered from Integrated DNA Technologies, and the full list can be found in Supplementary Table 2.

For step-by-step experimental protocol and details on reagents and concentrations, please refer to ^33^. Briefly, on the first day, sections were allowed to acclimate to room temperature, fixed with 3% paraformaldehyde (PFA), permeabilized with 0.1M HCL, dehydrated with ethanol, and incubated in reverse transcription master mix overnight at 37°C. Following reversed transcription, the sections were postfixed with 3% PFA and incubated in PLP mix for 30 min at 37°C and 90 min at 45°C. Then, sections were incubated in a rolling circle amplification mix for 4 hrs at 37°C and overnight at 30°C. On the third day, sections were washed with 2xSSC and incubated in BP mix for 1 hr at RT with light shaking. Following BP incubations, sections were washed with 2xSSC and incubated with DO mix (with Hoechst) for 1 hr at RT with light shaking. Sections were then washed with PBS, and a cover slip was applied using SlowFade gold antifade mounting medium. Samples were then imaged using a Leica DMi8 epifluorescence microscope equipped with LED light source (Lumencor SPECTRA X, nIR, 90-10172), sCMOS camera (Leica DFC9000 GTC, 11547007), and 20x objective (HC PC APO, NA 0.8, air). After imaging, sections were incubated in a stripping buffer for 30 min to strip off the DO and BP. This cycle of imaging, stripping, and incubation was repeated 4 more times using different BP oligos.

### HybISS images preprocessing

Each feld of view (FOV) was projected to obtain a flattened two-dimensional image using a variant of the extended-depth-of-field algorithm ^49^. Each FOV was then stitched using ASHLAR ^50^ and each hybridization cycle image was aligned to the first cycle via wsireg ^51^. HybISS signals (corresponding to diffraction-limited spots) were detected with Spotiflow ^52^ by applying the pre-trained *hybiss* model independently in each cycle and channel. Detections were assigned a gene identity (decoded) using the nearest-neighbor decoder implemented in starfish ^53^ using Spotiflow’s output probabilities as well as the spot intensities. Nuclei were segmented from the DAPI image of the first cycle with StarDist ^54^ using the 2D fluorescence versatile model. Nuclei masks were expanded for up to 20 µm each in order to retrieve cell masks, from which a cell-by-gene matrix was generated. Samples with detected genes lower than 180 were excluded from further analysis. Gene expressions were normalized and logarithmized using Scanpy ^55^. Differentially expressed genes (DEGs) across conditions were calculated using scanpy.tl.rank_genes_groups (Figure 4J and Figure S12, Supplementary Table 3).

### Morphological midline identification

The morphological midline of each budoid was identified using a custom Python tool (https://github.com/AztekinLab/MidlineIdentifier). The first step was to obtain a segmentation of the budoid structure. The binarised cell segmentation image of each budoid was subjected to image closing, hole filling and then opening. A disc footprint of radius 50 was used for closing and 20 for opening. For budoids located less than 50 pixels from the image boundary, an additional 100 pixels were padded before segmentation. All segments were identified using skimage.measure.label and skimage.measure.regionprops but only the largest segment was kept for structural segmentation. The remaining regions were considered as noise and were removed by setting the corresponding pixels to false. Next, the Euclidean distance transform matrix was computed on the segmentation using the scipy.ndimage.distance_transform_edt function, on which ridge detection was performed using skimage.filters.meijering with black_ridges set to False. To determine the two ends of the midline, the corners of the segmentation were detected using skimage.feature.corner_harris. The two corners closest to the two ends of the major axis based on Euclidean distance were defined as the ends of the midline to ensure that the morphological midline followed the direction of the major axis of the segmentation. Finally, using the difference between one and the ridge matrix as a scoring system, the midline was calculated by walking through the two corners using skimage.graph.route_through_array. Diagonal moves were allowed.

### Cell projection

Cells that fall out of the structure segmentation were removed. Simply projecting cells onto the nearest coordinate on the morphological midline leads to an overcrowdness problem where the majority of cells are associated with the same coordinate. Thus, we developed a scoring scheme which takes into account the distance between coordinates and cells and the number of cells associated with the coordinates. The score of the coordinate-cell pair is defined as

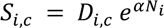

where represents the Euclidian distance and is the number of cells associated with it. The scaling factor is set to 0.01. Each cell was then projected to the coordinate with the highest score.

### Morphological midline orientation and gene quantification

To perform cross-sample comparisons, all the budoids need to be ordered in the same orientation. For this, we aimed to determine the proximal and distal domains of the morphological midline in each budoid. The two domains were determined using a set of chondrogenic (*Sox9, Acan, Col2a1, Col9a1, Col9a2, Col11a1*) and fibroblast genes (*Col1a1, Col3a1*). Budoids were first divided into four equidistant bins along the midline, within which gene expressions were averaged following scanpy.tl.score_genes to calculate the enrichment score of the proximal and distal genes, respectively. The half with a higher chondrogenic score was determined as the proximal. Midlines were then scaled to the range of zero to one to make it comparable across different budoids where zero represents the proximal, and one represents the distal. For line plots in Figure S14B, *statsmodels.ZeroInflatedPoisson* was used to fit the raw expression of each condition.

The proximal and distal scores were further used to classify the polarization property of budoids. Budoids with both proximal score and distal score greater than 0.01 were considered as polarized; budoids with either proximal score or distal score less than 0.01 were considered as non-polarized; budoids with both proximal and distal scores less than 0.01 were considered invalid samples. Polarized budoids can be divided into two equal parts from the middle along the morphological midline, the proximal and the distal.

### Principal component analysis (PCA)

Gene expression of each budoid or each part (proximal or distal) of the budoid was averaged. Genes expressed in less than 1% of cells were excluded. Missing values were imputed using the mean. The averaged expression matrix was standardized using sklearn.preprocessing.StandardScaler, followed by PCA using sklearn.decomposition.PCA (Figure 4K).

### Identification of the most representative section

For different sections from the same budoid, we remove the outlier sections based on the PCA. If these sections show different phenotypes (e.g., one polarizes but not the other), we remove all the sections. If these sections are similar, we 1) keep the middle section; 2) If there are sections from other budoids in the same slice that have already been retained, keep the section of similar morphological positions as the retained ones; 3) keep the morphologically more intact ones.

### Statistics and replicate information

Sample sizes were not pre-determined, and blinding was not applied during analyses. One-way ANOVA was used in Figure S1E, L, Figure S9D, E and Figure S11A, B, and t-test was used for Figure 4G, Figure S10B. Other performed statistical tests were indicated in the methods in relevant sections. Replicate information for experiments is provided in text or legends. Throughout the manuscript, n denotes technical replicates, and N denotes a biological replicate that is performed on a separate day, with a separate material, or both. All representative images are selected from experiments with at least n=3 and N=3. The standard error mean was provided for relevant data with (±standard error mean).

## Supplemental Figures

**Figure S1:**
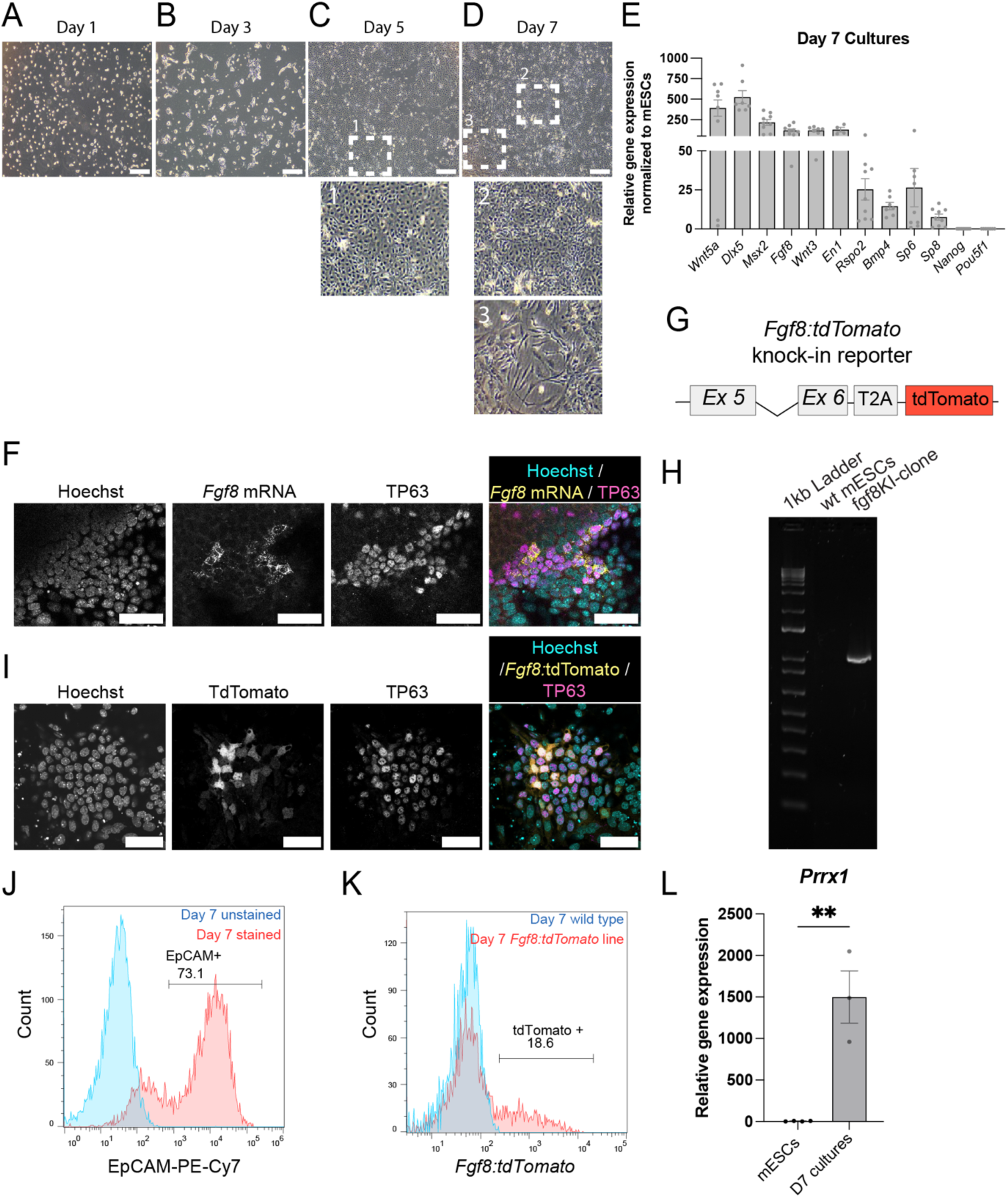
Characterization of stem-cell-derived heterogeneous cultures. (A-D) Representative brightfield images of induced cultures on Day 1 (A), Day 3 (B), Day 5 (C), and Day 7 (D). Insets highlight the homogeneous induction observed on Day 5 and the mixture of epithelial and mesenchymal cells at Day 7. Scale bar = 50 μm. (E) Quantitative RT-PCR results of the apical ectodermal ridge (AER) and pluripotent stem cell markers for Day 7 cultures. All samples were normalized to mouse embryonic stem cells (mESCs) and RPL27 housekeeping gene expression. Each dot represents the average of at least two technical replicates in a biological replicate. N>7 for all analyzed genes. (F) Representative confocal image of a cluster of *Fgf8* mRNA and TP63-positive AER-like cells. Cyan: Hoechst, Yellow: Fgf8 mRNA, magenta: TP63. Scale bar = 50 μm. (G) Schematic describing the knock-in to generate *the Fgf8:tdTomato* reporter line. (H) Genotyping result for the generated *Fgf8* reporter line, with primers targeting Exon 6 and *tdTomato*. (I) Representative confocal image of a cluster of *Fgf8:tdTomato* and TP63-positive AER-like cells. Cyan: Hoechst, Yellow: *Fgf8:tdTomato*, magenta: TP63. Scale bar = 50 m. (J) An example of flow cytometry data used to quantify EpCAM-positive cells in Day 7 cultures, leading to the Figure 1E, is shown. Unstained samples were used to distinguish the signal. (K) An example of flow cytometry data to quantify Fgf8:tdTomato positive cells in Day 7 cultures, leading to the Figure 1F, is shown. Wild-type controls were used to distinguish the signal. (L) Quantitative RT-PCR results of *Prrx1* expression in Day 7 (D7) cultures. All samples are normalized to RPL27 housekeeping gene expression. Each dot represents the average of at least two technical replicates in a biological replicate. N=4 for mESCs, and N=3 for Day 7 cultures.

**Figure S2:**
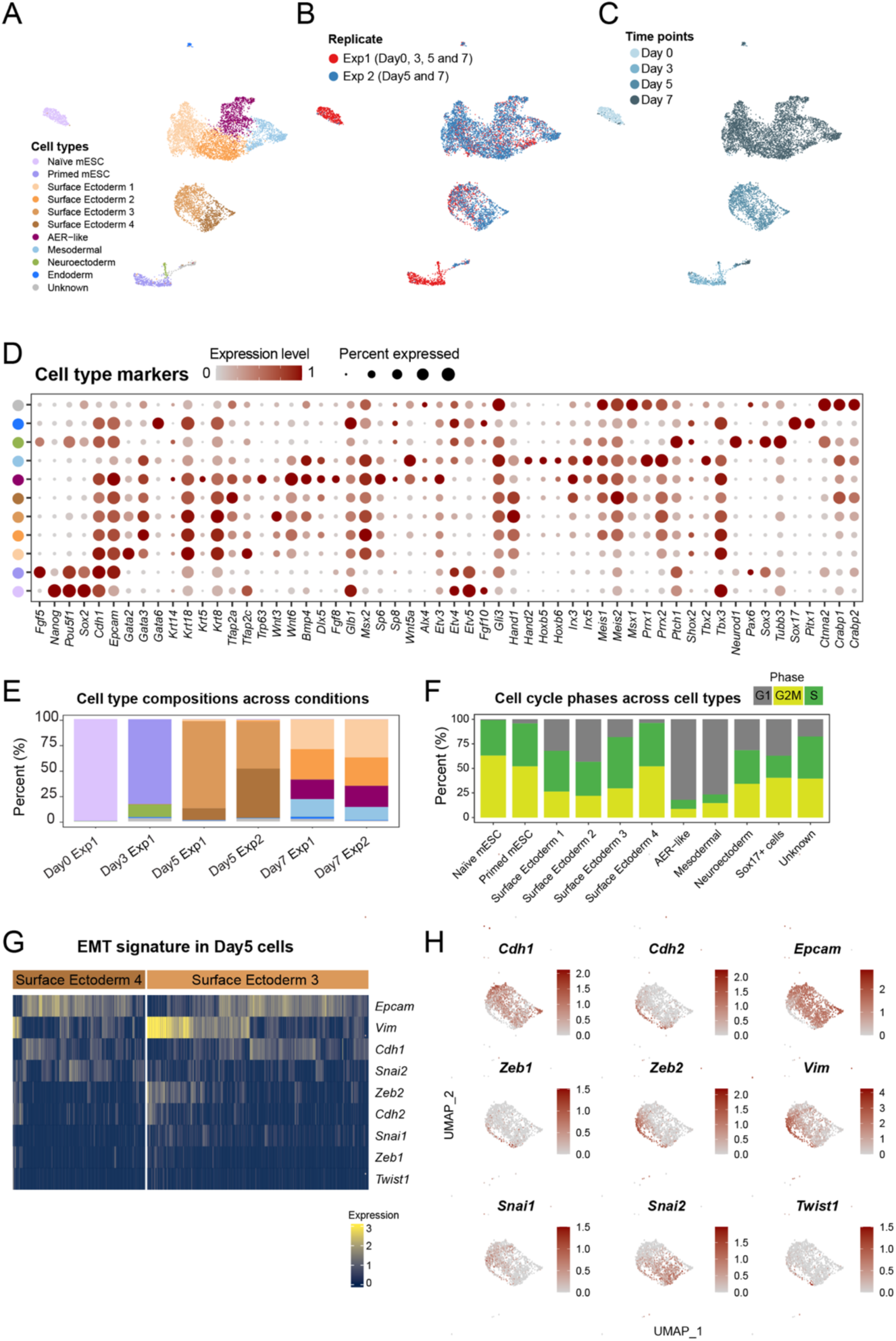
Single-cell RNA sequencing-based characterization of heterogeneous cultures. (A) UMAP representation of single-cell RNA sequencing (scRNA-Seq) displaying processed samples across different time points of the induction protocol. (B) UMAP representation of replicates. Replicate 1 contains cells from day 0, 3, 5, and 7 of the protocol, represented by red cells. Replicate 2 contains cells from day 5 and 7 of the protocol, represented by blue cells. (C) UMAP representation of cells from different time points in the dataset. Shades of light to dark blue indicate day 0 to day 7 samples. (D) Dot plot showing marker expressions used to annotate cell types. The size of the dot represents the percentage of cells expressing each marker. (E) Proportion analysis showing the relative abundance of each annotated cell type within the culture at each time point and replicate. Please note that some of these data are the same as in Fig 1H. (F) ScRNA-Seq-based cell cycle analysis for detected clusters. (G) Heatmap showing epithelial-to-mesenchymal transition (EMT) associated gene expression in day 5 surface ectoderm clusters. Each column represents one cell. (H) UMAP showing EMT-associated gene expression in day 5 surface ectoderm clusters. Shades of red indicate the expression level.

**Figure S3:**
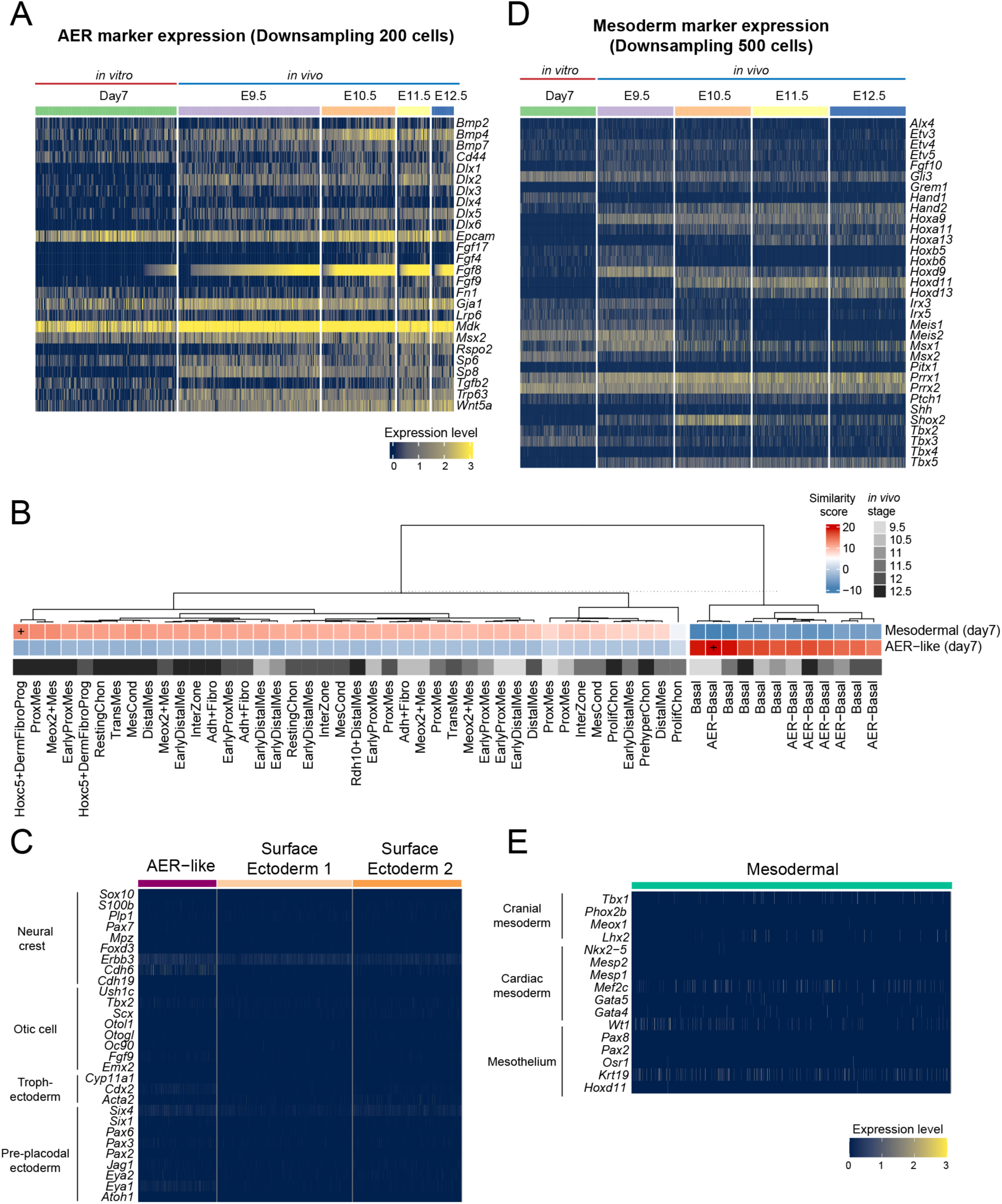
Heterogeneous cultures show varying degrees of similarity to in vivo limb cells. (A) Heatmap showing AER-associated gene expression levels in the generated *in vitro* AER-like cluster and *in vivo* AER clusters at developmental stages E9.5 to E12.5. *In vivo* data are obtained from Allou et al. (E9.5) ^56^ and Desanlis et al. (E10.5-12.5) ^57^. Each column represents one cell. (B) Heatmap displaying the transcriptome-wide similarity of generated in vitro AER-like and mesodermal cells at day 7 with in vivo limb cell types across stages E9.5 to E12.5. A ‘+’ symbol denotes the highest level of similarity of each row, and varying shades of gray at the bottom of the heatmap indicate the *in vivo* developmental stages. *In vivo* data and cluster annotations are obtained from Zhang et al.^22^. (C) Heatmap illustrating the expression levels of relevant ectodermal cell type markers in the generated *in vitro* AER-like cluster and surface ectoderm clusters in day 7 cultures. Each column represents one cell. (D) Heatmap presenting the expression levels of early lateral plate mesoderm and limb bud mesoderm markers in the generated in vitro mesoderm cluster compared with in vivo limb bud mesoderm clusters at stages E9.5 to E12.5. *In vivo* data and cluster annotations are obtained from Allou et al. (E9.5) ^56^ and Desanlis et al. (E10.5-12.5) ^57^. Each column represents one cell. (E) Heatmap showing relevant mesodermal cell type marker expression levels in the generated *in vitro* mesodermal cluster at day 7 cultures.

**Figure S4:**
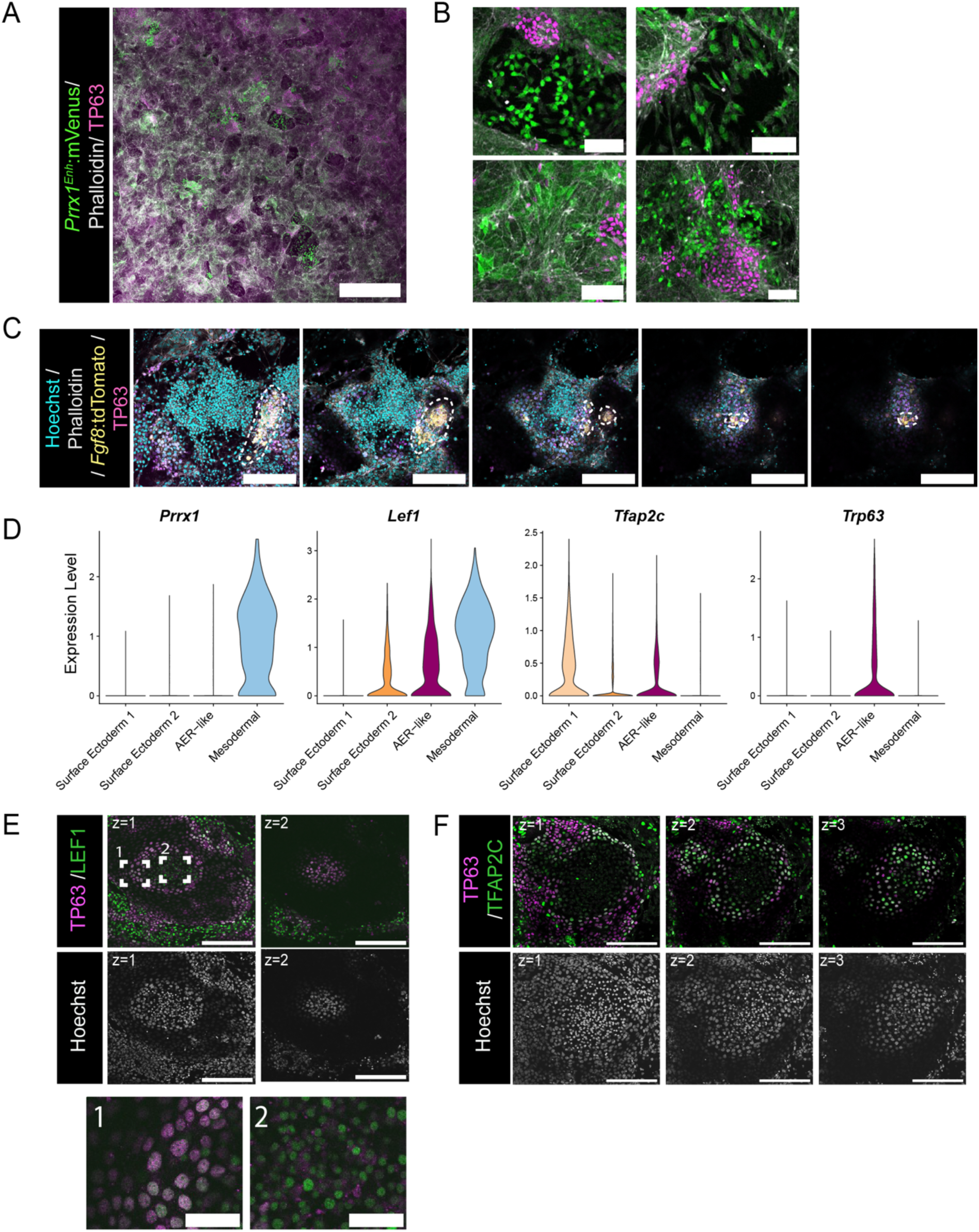
Generated epithelial and mesenchymal cells self-organize into domes. (A) Representative confocal image showing the distribution of *Prrx1^Enh^:mVenus* and TP63 positive cells in day 7 cultures. Green: *Prrx1^Enh^:mVenus*, Gray: Phalloidin, Magenta: TP63; Scale bar = 1000 μm. (B) Higher magnification images of *Prrx1^Enh^:mVenus* and TP63-positive cells within day 7 cultures. Note the fibroblast-like and circular morphologies of the *Prrx1^Enh^:mVenus* positive cells. Green: *Prrx1^Enh^:mVenus*; White: Phalloidin; Magenta: TP63. Scale bar = 100 μm. (C) Sequential Z-stack confocal images of a dome. The left to right panels show progressing depth through the Z-stack. Note that *Fgf8:tdTomato*-positive cells are located at the corners and tip of the dome and are circled with a dashed line. Z-stacks are denoted by z numbers. Cyan: Hoechst; White: Phalloidin; Yellow: *Fgf8:tdTomato*; Magenta: TP63. Scale bar = 200 μm. (D) Violin plots illustrating *Prrx1*, *Lef1*, *Tfap2c*, and *Tp63* expression levels across the main identified clusters in the day 7 scRNA-Seq dataset. Note that *Lef1* positive/*Tfap2c* negative/*Trp63* negative cells only indicate the mesodermal cluster. (E) Sequential Z-stack confocal images of a dome. Left to right panels represent progressing depth through the Z-stack, denoted by z numbers. Note that the larger nuclei of TP63-positive cells form the outer layer compared to LEF1-positive cells at the core of the dome. Gray: Hoechst; Green: LEF1; Magenta: TP63. Scale bar = 200 μm. Insets 1 and 2 offer higher magnification views of selected areas. Scale = 50 μm. (F) Sequential Z-stack confocal images of a dome structure. Left to right panels represent progressing depth through the Z-stack, denoted by z numbers. Green: TFAP2C; Magenta: TP63; Gray: Hoechst. Scale bar = 200 μm.

**Figure S5:**
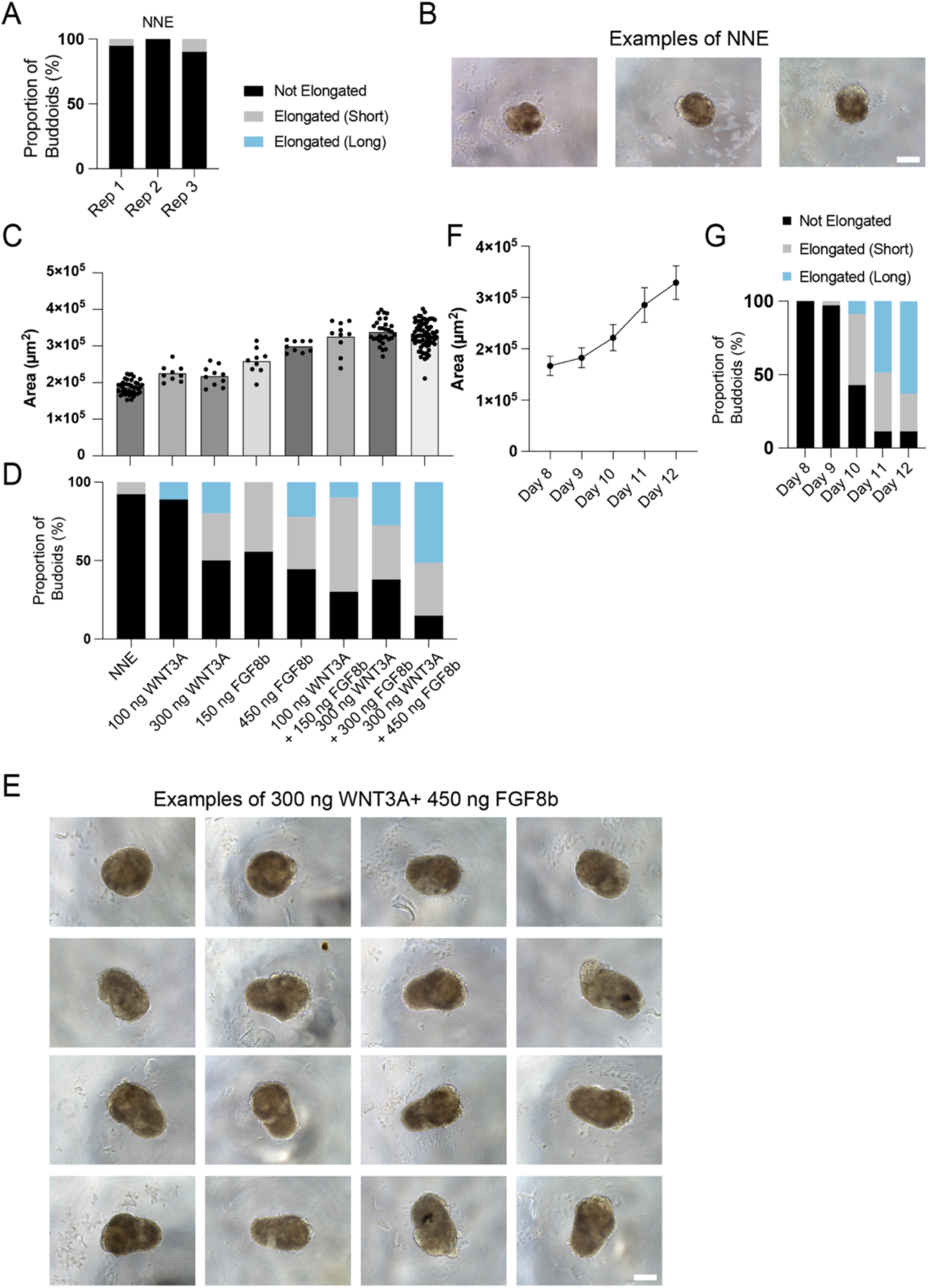
FGF8b and WNT3a treatment enhance the efficiency of budoid formation from heterogenous cultures-derived aggregates. (A) Bar chart illustrating the proportion of budoids elongating in no treatment condition at day 12. The total number of aggregates analyzed was n = 39 and N = 3. (B) Brightfield images of no treatment condition on day 12. Scale bar = 100 μm. (C-D) (C)Bar plot demonstrating the day 12 budoids area resulting from various FGF8b and WNT3a treatment concentrations. (D) Bar chart displaying the proportion of budoids exhibiting no elongation, elongation (short), or elongation (long) after treatment with different concentrations of growth factors. NNE is a treatment condition. NNE: n= 39, N=3: 100 ng/ul WNT3A: n= 9, N=1: 300 ng/ul WNT3A n=10, N=1; 150 ng/ul FGF8b, n=9, N=1: 450 ng/ul FGF8b n=9, N=1: 100 ng/ul WNT3A and 150 ng/ul FGF8b, n=10, N=1: 300 ng/ul WNT3A and 300 ng/ul FGF8b, n=30, N=2; 300 ng/ul WNT3A and 450 ng/ul FGF8b, n=72, N=2. Please note that some of these data are the same as in Fig 3B. (E) Examples of brightfield images showing day 12 budoids with varying elongation phenotypes after treatment with 300 ng WNT3A and 450 ng FGF8b. Scale bar = 100 μm. (F-G) (F) Line plot depicting the change in area from day 8 to day 12 of the protocol for budoids under the influence of FGF8b and WNT3a treatment. (G) Bar chart presenting the elongation phenotypes of budoids over time. Total aggregates analyzed: n = 94 for days 8, 9, 11, and 12; and n= 45 for day 10, N=2.

**Figure S6:**
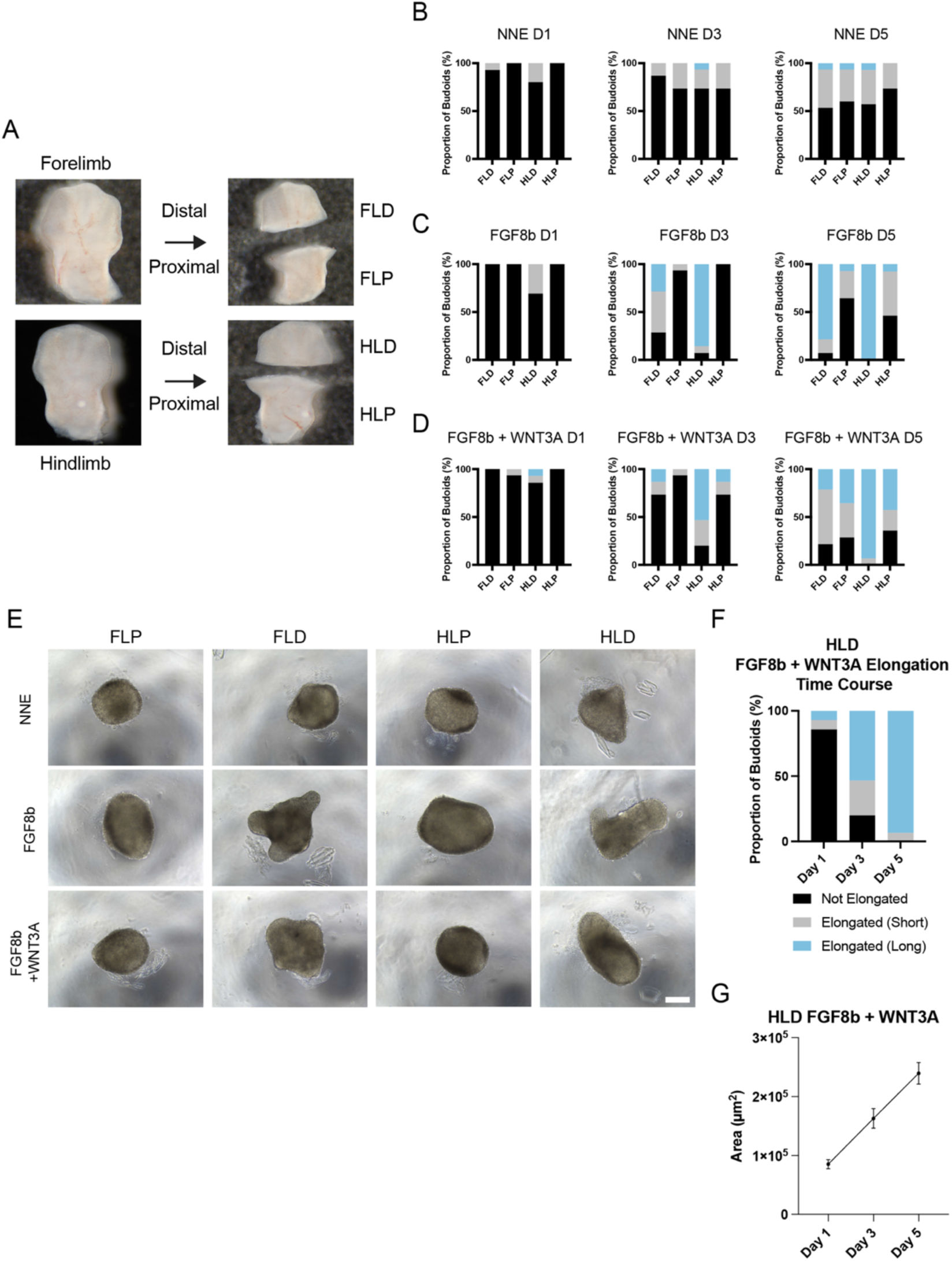
FGF8b and WNT3a treatment enhance the formation efficiency of in vivo-derived budoids. (A) Images of dissected forelimbs and hindlimbs from E12.5 mouse embryos used to generate budoids. The forelimb distal (FLD), forelimb proximal (FLP), hindlimb distal (HLD), and hindlimb proximal (HLP) regions are indicated. (B-D) Bar charts showing the proportion of budoids exhibiting various elongation phenotypes on Day 1, Day 3, and Day 5 after treatments with NNE, FGF8b, and a combination of FGF8b with WNT3a. The total number of budoids analyzed was n >14 for all samples, N = 3. (E) Brightfield images of Day 5 budoids displaying varying elongation phenotypes after treatments with NNE, FGF8b, and a combination of FGF8b with WNT3a across different collected limb samples. Scale bar = 100 μm. (F) Bar chart illustrating the time-course elongation phenotypes of HLD budoids treated with FGF8b and WNT3a. The total number of budoids analyzed n = 15, N = 3. (G) Line plot showing the time course area of HLD budoids treated with FGF8b and WNT3a. The total number of budoids analyzed n =15, N = 3.

**Figure S7.**
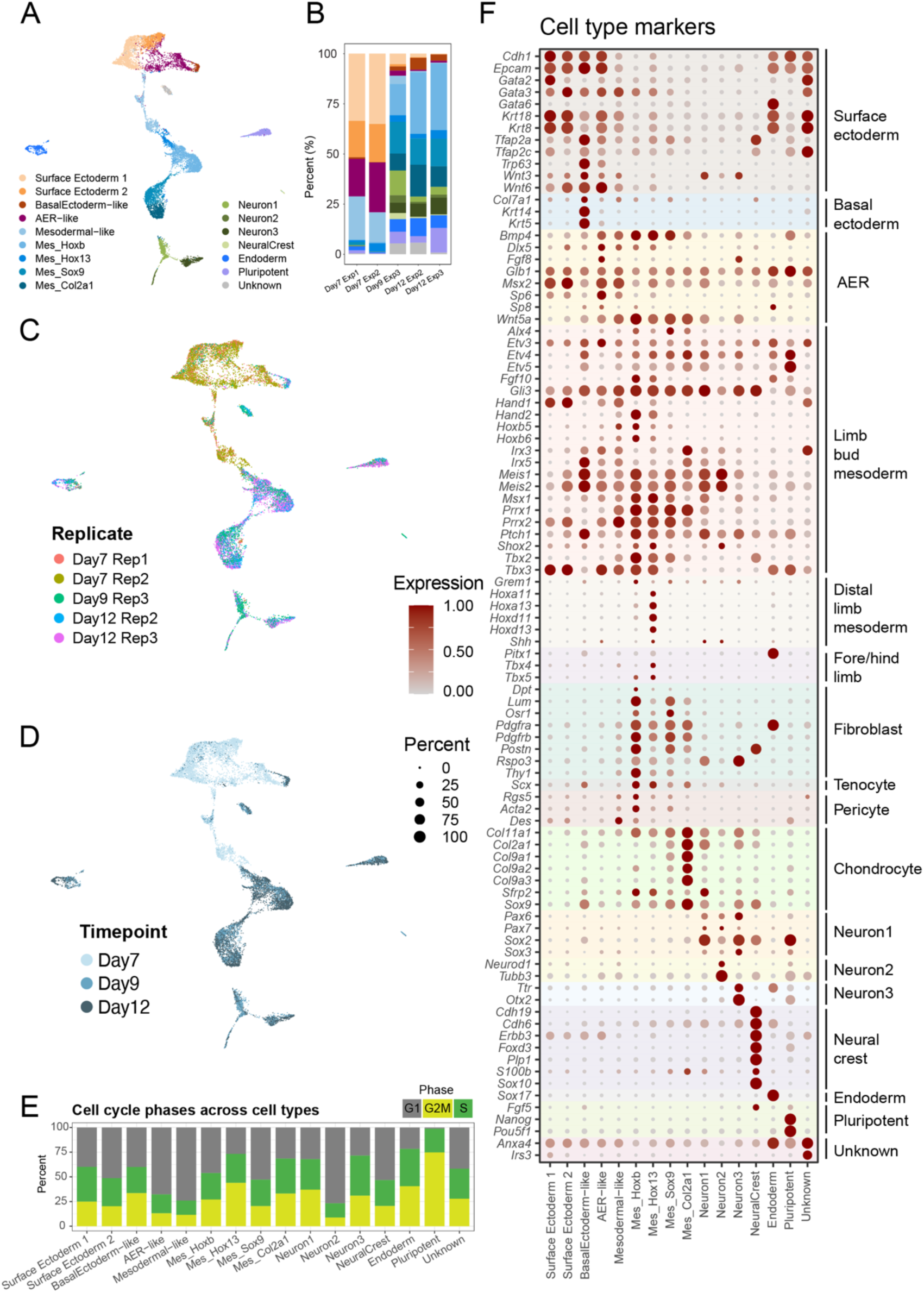
Single-cell RNA sequencing-based characterization of budoids. (A) UMAP visualization of scRNA-Seq data from budoids at different culture time points, as highlighted in Fig 3A, with each cell type uniquely colored. (B) Bar chart representation of the relative proportions of the cell types identified in the scRNA-Seq analysis. (C) UMAP plot showing the distribution of cells from different replicates. (D) Temporal UMAP representation of cells color-coded by time point from day 7 (light blue) to day 12 (dark blue) samples. (E) Bar chart summarizing the cell cycle phase distribution across all detected cell populations. (F) Dot plot characterizing the expression of marker genes that define each cell type. Dot size corresponds to the percentage of cells expressing a particular gene, while color intensity reflects expression level. Please note that the genes annotated with specific cell identities are not exclusive and can be expressed across different cell types.

**Figure S8:**
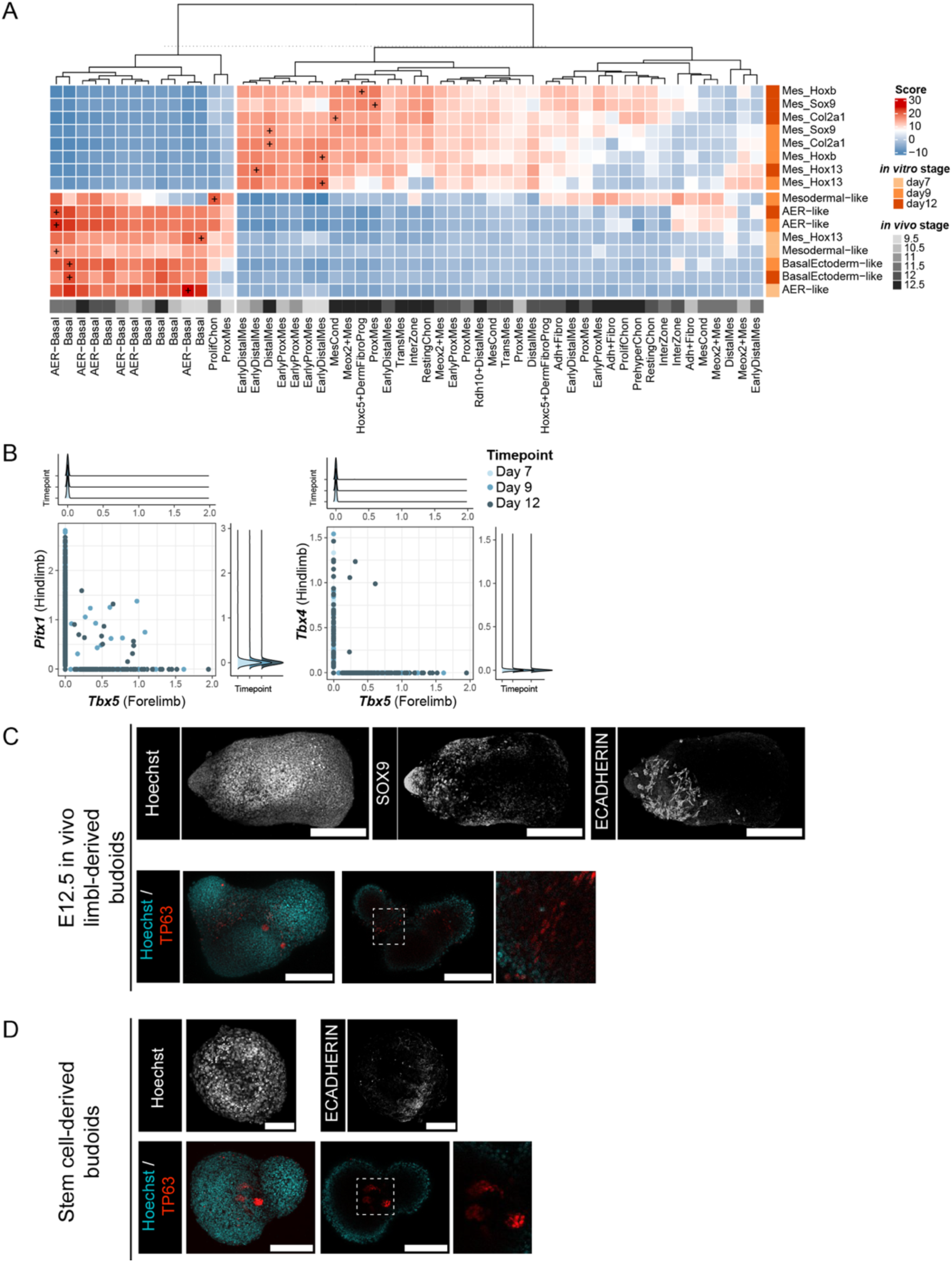
Stem cell-derived budoids show similarities to developing limbs *in vivo*, and ectodermal cells are internalized in stem cell or *in vivo*-derived budoids. (A) Heatmap presenting the transcriptome-level similarities between *in vitro*-generated AER-like and mesodermal cells during budoids generation over days 7 to 12, and *in vivo* limb cell types at developmental stages E9.5 to E12.5. ‘+’ indicates the highest similarity in each row, while the gray gradient at the bottom of the heatmap reflects the developmental stages. *In vivo* data and cluster annotations are obtained from Zhang et al. ^22^. (B) Scatter plots demonstrating expression patterns of hind and fore limb identity-related genes in individual cells across various time points. Each dot represents a single cell, and time points are distinguished by color shading. (C) (Top) Max projection confocal image showing day 5 in vivo-derived budoids with polarized SOX9 expression and limited E-CADHERIN. Scale bar = 200 μm. (Bottom-left) Max projection confocal image showing TP63-positive cells. (Bottom-mid) An optical section of the budoid shows TP63-positive cells internally localized within the structure. Scale bar = 200 μm. (Bottom-right) Zoomed-in view of the optical section. (D) (Top) Max projection confocal image showing day 9 in stem cell-derived budoids with limited E-CADHERIN. Scale bar = 200 μm. (Bottom-left) Max projection confocal image showing day 12 budoid with TP63 positive cells. (Bottom-mid) An optical section of the budoid shows TP63-positive cells internally localized within the structure. Scale bar = 200 μm. (Bottom-right) Zoomed-in view of the optical section.

**Figure S9.**
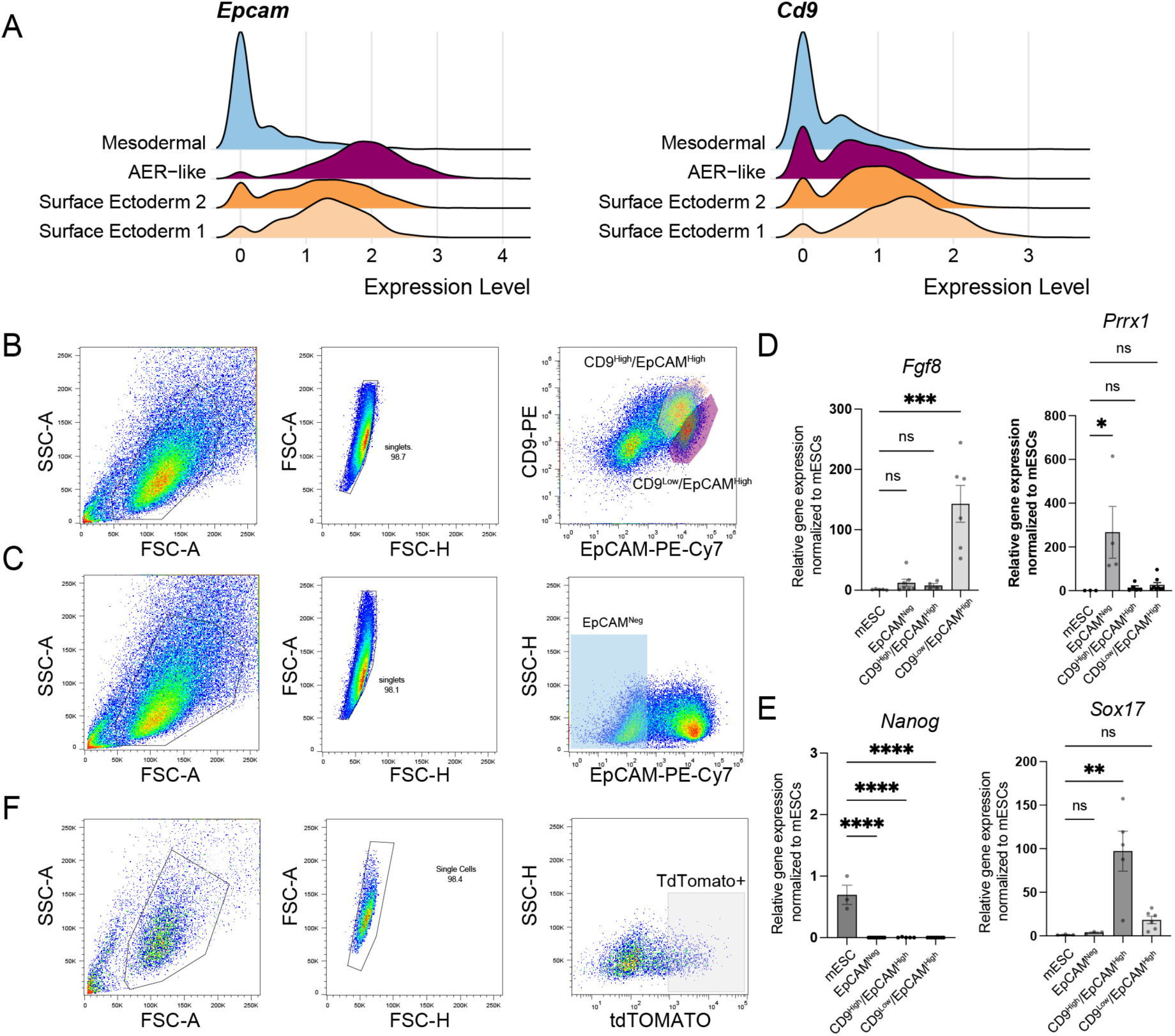
Cell sorting strategy to enrich mesodermal, AER-like, and surface-ectoderm-like cells. (A) Histogram plots illustrate the expression of *Epcam* and *Cd9* in Day 7 clusters identified by scRNA-seq data. (B) Representative flow cytometry data showing antibody-based labeling with CD9 and EpCAM generates three populations. An enrichment strategy based on this is outlined for AER-like (CD9^Low^/EPCAM^High^) and surface-ectoderm-like (CD9^High^/EPCAM^High^) populations. FACS-enriched populations are shown with shaded red for AER-like cells and yellow for surface ectoderm-like cells. (C) Representative flow cytometry data showing antibody-based labeling with EpCAM for isolating mesodermal cells (EpCAM^Neg^). The FACS-enriched population is shown in shaded blue. (D) Quantitative RT-PCR results for (left) *Fgf8* and (right) *Prrx1* in the FACS-enriched populations against mouse embryonic stem cells (mESCs). All samples are normalized to RPL27 housekeeping gene expression. Each dot represents the average of at least 2 technical replicates for one biological replicate. N >3 for all conditions. (E) Quantitative RT-PCR results for (left) *Nanog* and (right) *Sox17* in the FACS-enriched populations against mESCs. All samples are normalized to RPL27 housekeeping gene expression. Each dot represents the average of at least 2 technical replicates for one biological replicate. N >3 for all conditions. (F) Representative flow cytometry data showing the enrichment strategy for isolating AER-like cells using the *Fgf8:tdTomato* reporter line. The FACS-enriched population is shown with shaded gray.

**Figure S10.**
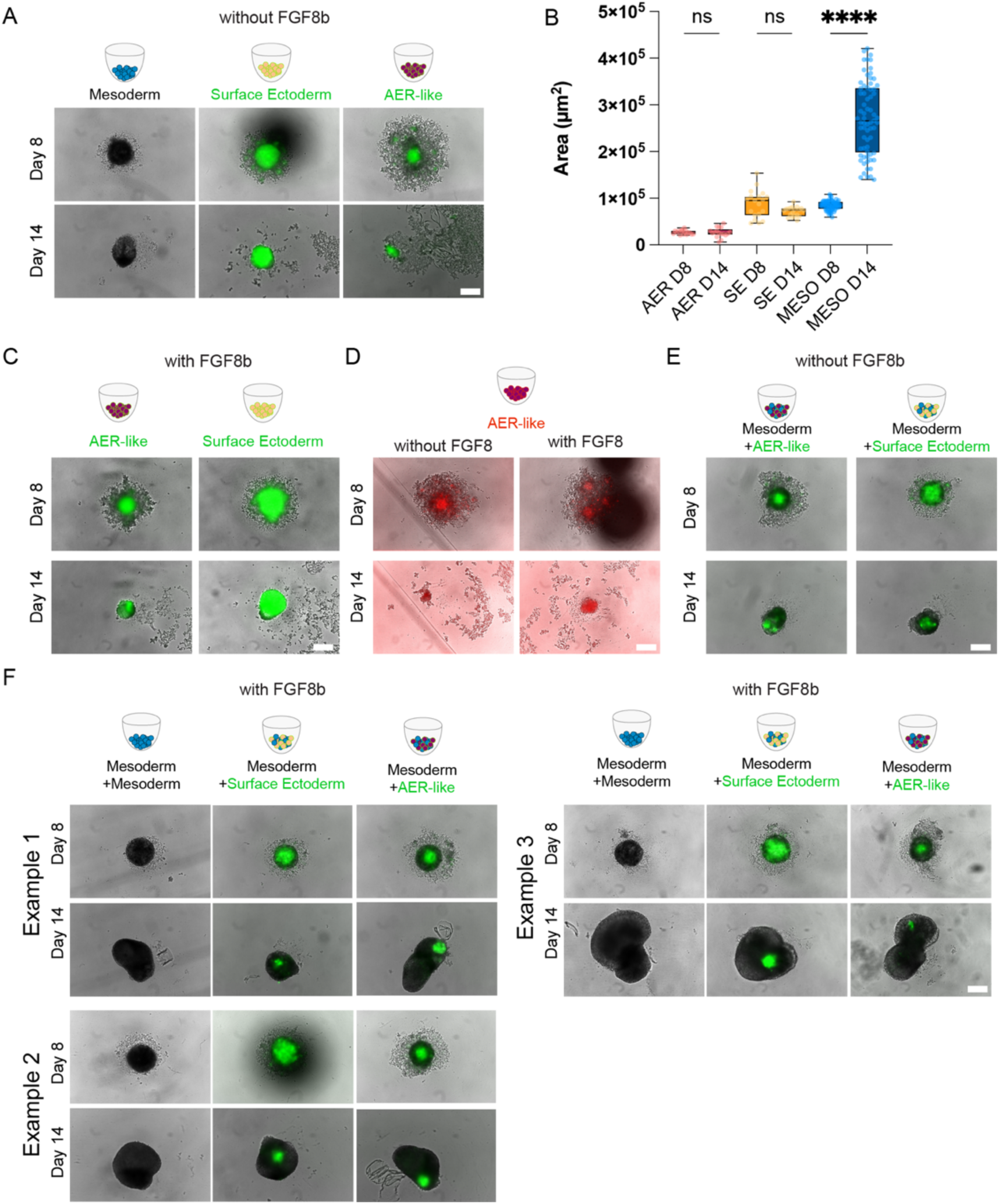
FGF8b treatment is necessary for budoids generation. (A) Representative images display FACS-enriched EpCAM^Neg^ mesodermal, CD9^High^/EpCAM^High^ surface ectoderm-like, CD9^Low^/EpCAM^High^ AER-like populations without FGF8b treatment at Day 8 and Day 14 of the protocol. Note excessive non-aggregated and shed cells, especially in ectodermal cells. Scale bar = 100 μm. (B) A box plot indicates the changes in area measurements for the FACS-enriched populations on Day 8 and Day 14 of the protocol. D8 denotes day 8 of the protocol, and D14 denotes day 14 of the protocol. Total number of aggregates from AER-like n= 17 from N=4, SE-like= 19 from N=4, and mesodermal n= 69, from N= 9, n.s.=not significant; **** denotes p= <0.0001; (C) Representative images showing FACS-enriched CD9^Low^/EpCAM^High^ AER-like and CD9^High^/EpCAM^High^ surface ectoderm-like cells with FGF8b treatment at Day 8 and Day 14. Please note excessive non-aggregated and shed cells, particularly in day 8 ectodermal cells. Scale bar = 100 μm. (D) Representative images showing FACS-enriched *Fgf8:tdTomato* positive AER-like cells from *Fgf8:tdTomato* mESC line, with and without FGF8b treatment. Note excessive non-aggregated and shed cells. Scale bar = 100 μm. (E) Representative images of recombination of mesoderm with AER-like or surface ectoderm-like cells without FGF8b treatment at Day 8 and Day 14 of the protocol are shown. Note excessive non-aggregated and shed cells, especially in day 8 ectodermal cells. Scale bar = 100 μm. (F) Additional examples of recombinant budoids with FGF8b treatment at Day 8 and Day 14 of the protocol. These images show the range of phenotypes recorded, albeit lesser in frequency compared to those shown in Figure 4 C-E. Images in the same example group show structures from the same experiment. Note excessive non-aggregated and shed cells, especially in ectodermal cells. Scale bar = 100 μm.

**Figure S11.**
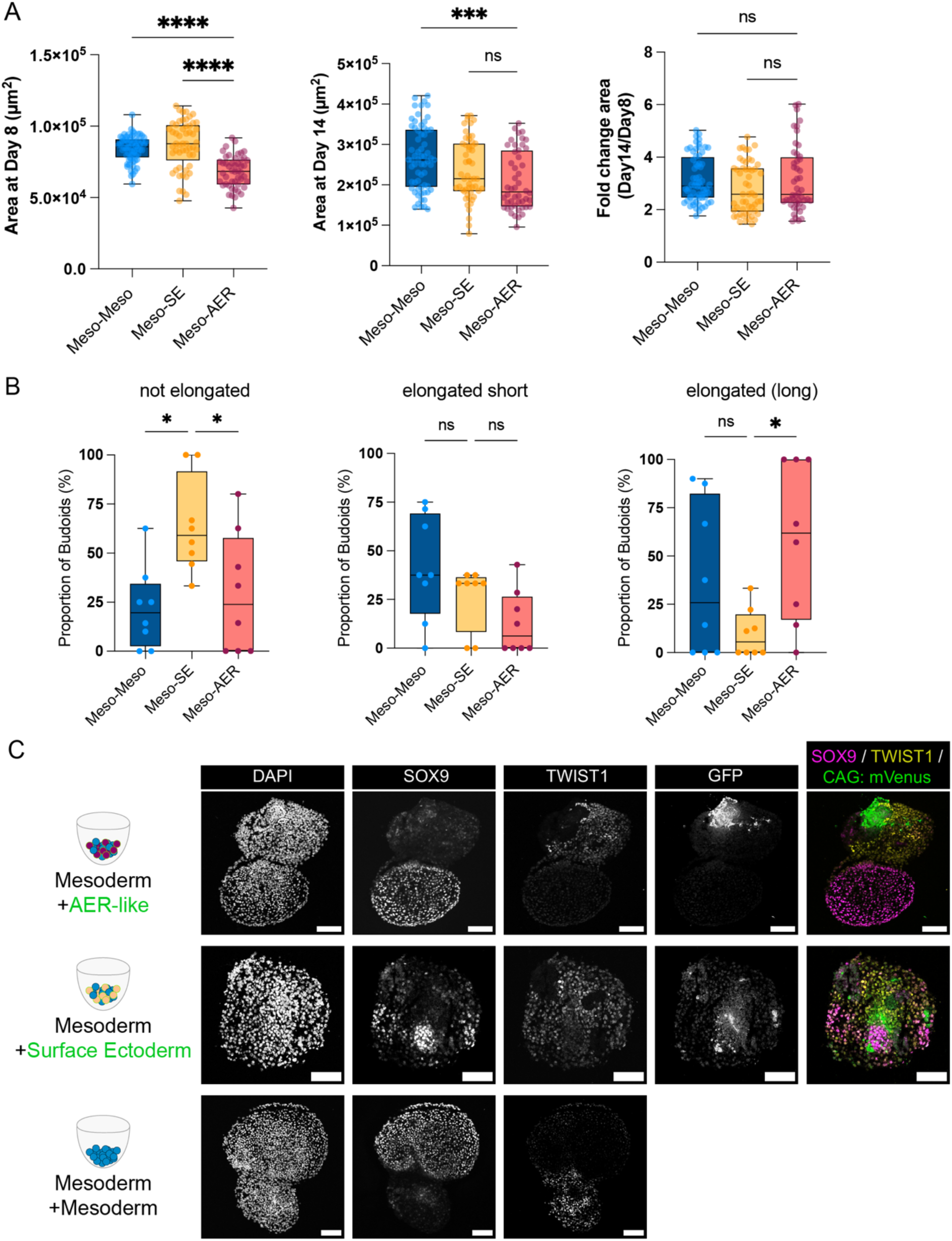
Recombinations show no significant difference in area fold change, but they influence spatial organization. (A) Box plots illustrating area measurements for FACS-enriched populations on (Left) day 8, (Mid) day 14, and (Right) the fold change in area from day 8 to day 14 for individual budoids. Each dot represents one recombinant budoid. The total number of budoids analyzed for Meso-Meso n= 64; Meso-SE n=53, Meso-AER n= 48, all from N=8. n.s.= not significant; *** denotes p< 0.0005, **** denotes p< 0.0001; (B) Box plots depicting the proportions of budoids exhibiting various elongation phenotypes in recombination experiments for (Left) non-elongated, (Middle) elongation (short), and (Right) elongation (long). Each dot represents the proportion of one biological replicate obtained from 3-10 technical replicates. N= 9. (C) Representative confocal images of sectioned recombinant budoids are provided to illustrate the spatial organization within these structures. Magenta: SOX9, Yellow: TWIST1, Green: GFP. Scale bar = 100 μm.

**Figure S12.**
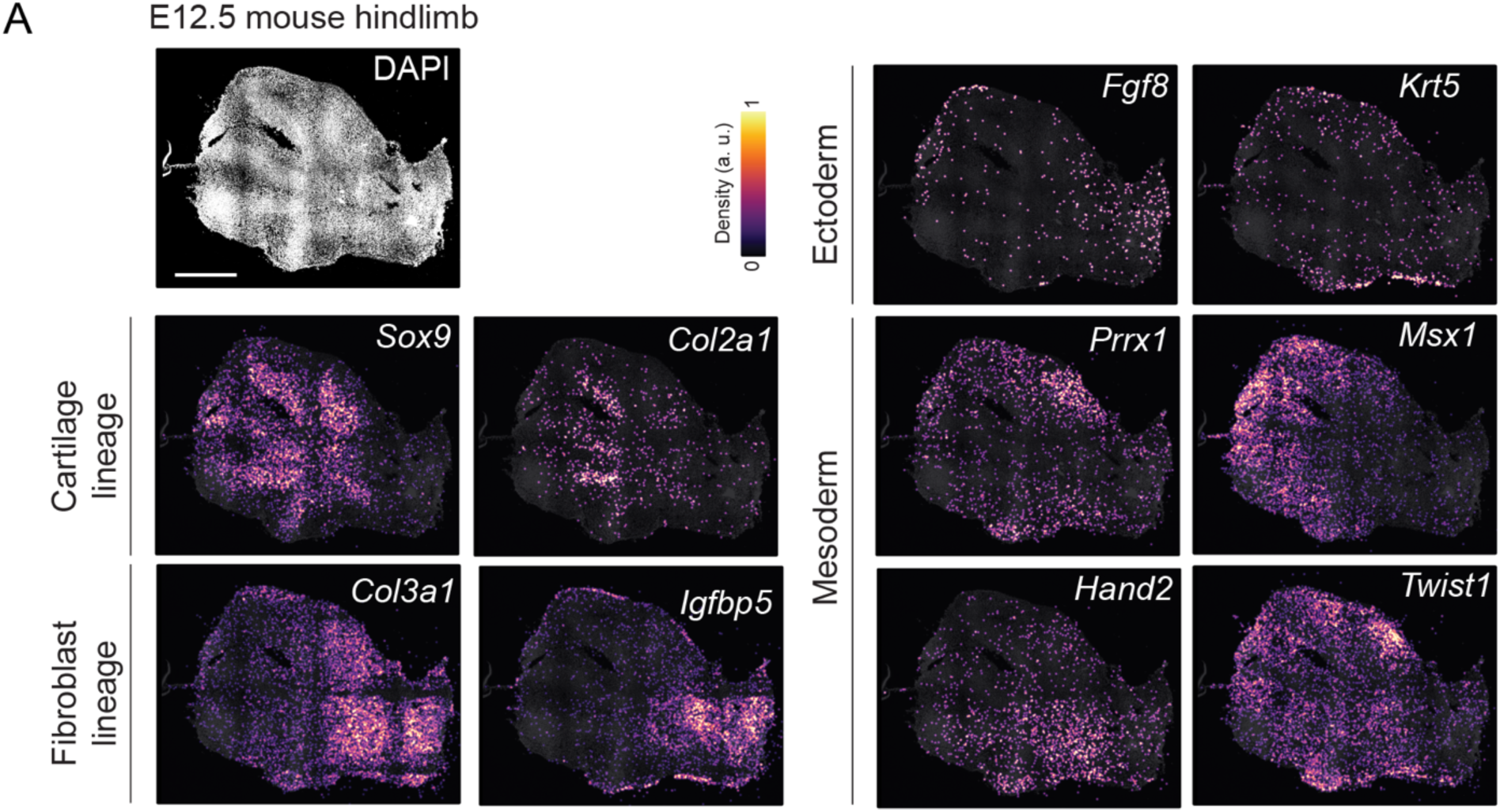
Hybridization-based in situ sequencing (HybISS) detects known limb developmental gene expression patterns. Example Hybridization-based in situ sequencing (HybISS) results for limb development-associated genes. Scale bar = 500 μm.

**Figure S13.**
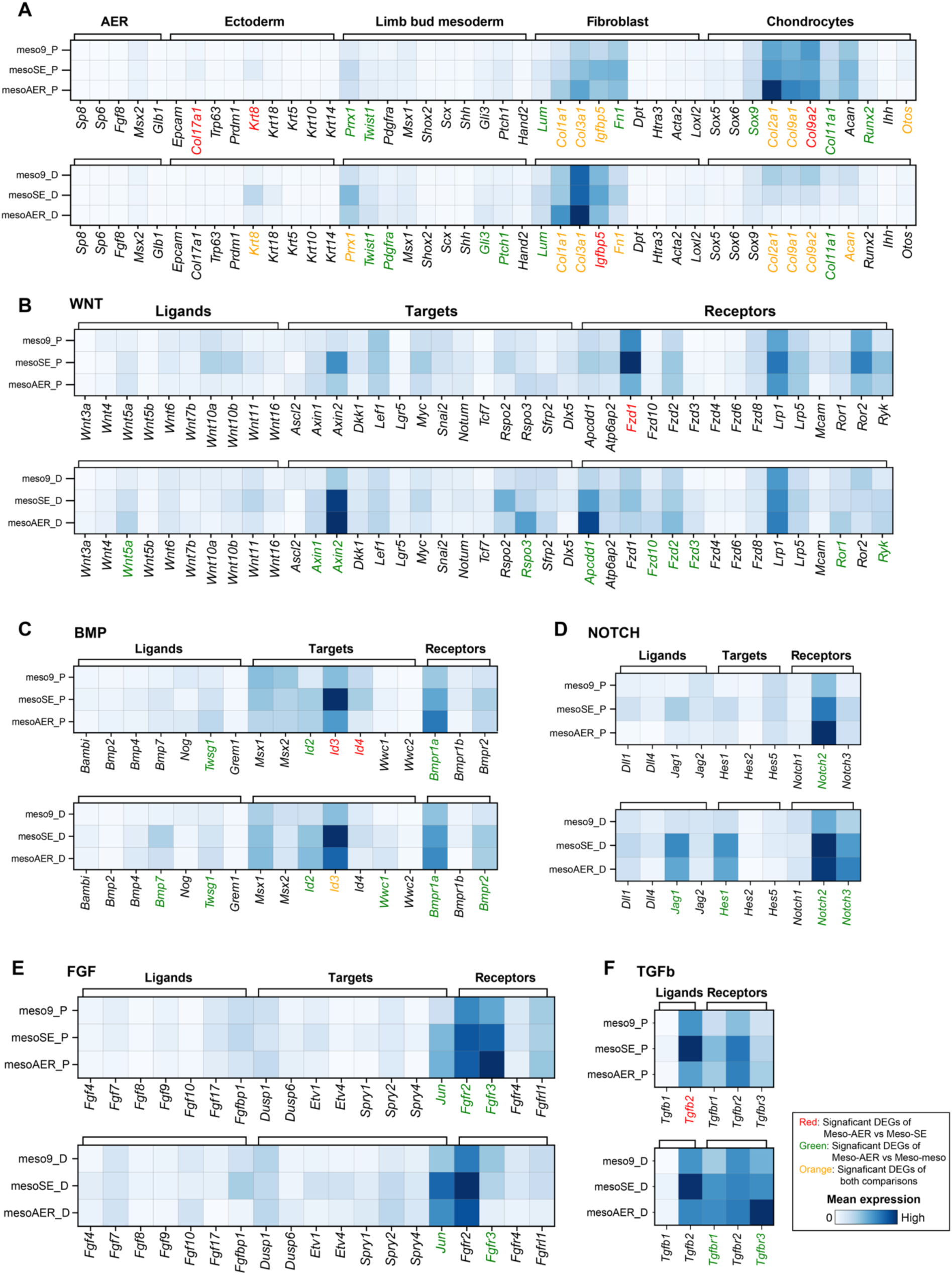
HybISS reveals recombination-induced molecular changes in proximal and distal domains. Heatmaps showing average expression levels of genes associated with (A) limb development-related cell fates, (B) WNT, (C) BMP, (D) NOTCH, (E) FGF, and (F) TGFb pathways in the recombinant budoids single cell distal and proximal domains. Genes with statistically significant differences (adjusted P-values < 0.05) are highlighted, with green indicating a significant difference between mesoderm-AER and mesoderm-mesoderm comparisons, red signifies significant differences between mesoderm-AER and mesoderm-surface ectoderm and orange indicating significance in both comparisons. Number of HybISS sections analyzed for Meso-Meso n= 20, N=4, Meso-SE n=12, N=3, Meso-AER n=20, N= 4.

**Figure S14.**
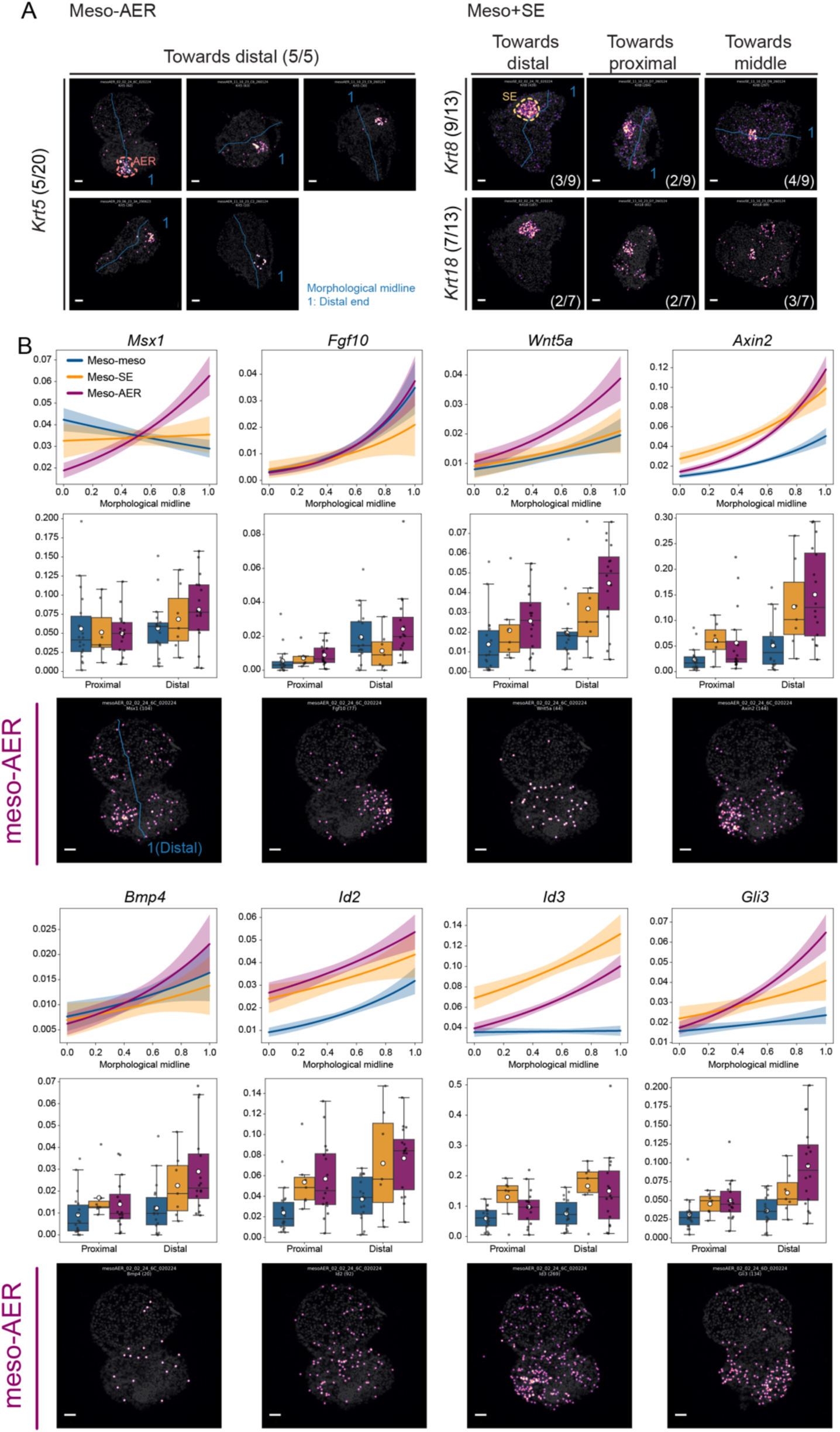
HybISS demonstrates the lasting impact of AER-like recombinations. (A) (Left) *Krt5* expression marks the remnants of AER-like cells in meso-AER recombinant sections, with a 5/20 prevalence. Notably, all 5/5 cases with *Krt5* positive cells are localized to the distal domain of the recombinants. (Right) In meso-SE recombinants, remnants of surface-ectoderm-like cells are identifiable through *Krt8* in 9/13 instances and *Krt18* in 7/13 instances. These markers do not exhibit preferential expression towards distal or proximal domains. Scale bar =50 μm. (B) Previously suggested AER induced *Msx1, Fgf10, Wnt5a, Axin2, Bmp4, Id2, Id3*, and *Gli3* in recombinant budoids are shown. For each gene: (Top) Line graphs plot the gene expression gradients from the proximal to distal ends of recombinant budoids. Each condition is color-coded. The zero-inflated Poisson model was used to fit the data in each condition. Confident intervals were shown in shades. (Middle) Box plots show the normalized expression levels of these genes within distal versus proximal domains across different conditions. (Bottom) Accompanying example images display HybISS results for the specified genes in meso-AER samples. Scale bar =50 μm.

## Supplementary Tables

Supplementary Table 1. Quality assessment and details of single-cell RNA-seq datasets.

Supplementary Table 2. Probes sequences of HybISS

Supplementary Table 3. Differentially expressed gene analysis of HybISS dataset

Supplementary Table 4. Primers used for cloning, PCR and qPCR

## References

1. Aztekin, C. Tissues and Cell Types of Appendage Regeneration: A Detailed Look at the Wound Epidermis and Its Specialized Forms. Frontiers in Physiology 12, 2047 (2021).

2. Fernandez-Teran, M. & Ros, M. A. The Apical Ectodermal Ridge: morphological aspects and signaling pathways. Int. J. Dev. Biol. 52, 857–871 (2008).

3. Zeller, R., López-Ríos, J. & Zuniga, A. Vertebrate limb bud development: moving towards integrative analysis of organogenesis. Nat Rev Genet 10, 845–858 (2009).

4. Geetha-Loganathan, P., Nimmagadda, S., Christ, B., Huang, R. & Scaal, M. Ectodermal Wnt6 is an early negative regulator of limb chondrogenesis in the chicken embryo. BMC Developmental Biology 10, 32 (2010).

5. Lee, B. H., Seijo-Barandiaran, I. & Grapin-Botton, A. Epithelial morphogenesis in organoids. Current Opinion in Genetics & Development 72, 30–37 (2022).

6. Zhao, Z. et al. Organoids. Nat Rev Methods Primers 2, 1–21 (2022).

7. Saunders, J. W. The proximo-distal sequence of origin of the parts of the chick wing and the role of the ectoderm. J Exp Zool 108, 363–403 (1948).

8. Summerbell, D. A quantitative analysis of the effect of excision of the AER from the chick limb-bud. J Embryol Exp Morphol 32, 651–660 (1974).

9. Chen, Y., Xu, H. & Lin, G. Generation of iPSC-derived limb progenitor-like cells for stimulating phalange regeneration in the adult mouse. Cell Discov 3, 17046 (2017).

10. Atsuta, Y. et al. Direct reprogramming of non-limb fibroblasts to cells with properties of limb progenitors. Dev Cell 59, 415–430.e8 (2024).

11. Smith, C. A. et al. Directed differentiation of hPSCs through a simplified lateral plate mesoderm protocol for generation of articular cartilage progenitors. PLOS ONE 18, e0280024 (2023).

12. Loh, K. M. et al. Mapping the pairwise choices leading from pluripotency to human bone, heart and other mesoderm cell-types. Cell 166, 451–467 (2016).

13. Mori, S. et al. Self-organized formation of developing appendages from murine pluripotent stem cells. Nat Commun 10, 3802 (2019).

14. Boutin, E. L. & Fallon, J. F. Insulin improves survival but does not maintain function of cultured chick wing bud apical ectodermal ridge. Anat Rec 233, 467–477 (1992).

15. Koehler, K. R., Mikosz, A. M., Molosh, A. I., Patel, D. & Hashino, E. Generation of inner ear sensory epithelia from pluripotent stem cells in 3D culture. Nature 500, 217–221 (2013).

16. Selever, J., Liu, W., Lu, M.-F., Behringer, R. R. & Martin, J. F. *Bmp4* in limb bud mesoderm regulates digit pattern by controlling AER development. Developmental Biology 276, 268–279 (2004).

17. Min, H. et al. Fgf-10 is required for both limb and lung development and exhibits striking functional similarity to Drosophila branchless. Genes Dev 12, 3156–3161 (1998).

18. Barrow, J. R. et al. Ectodermal Wnt3/β-catenin signaling is required for the establishment and maintenance of the apical ectodermal ridge. Genes Dev 17, 394–409 (2003).

19. Haro, E. et al. Sp6 and Sp8 Transcription Factors Control AER Formation and Dorsal-Ventral Patterning in Limb Development. PLOS Genetics 10, e1004468 (2014).

20. Storer, M. et al. Senescence Is a Developmental Mechanism that Contributes to Embryonic Growth and Patterning. Cell 155, 1119–1130 (2013).

21. Muñoz-Espín, D. et al. Programmed cell senescence during mammalian embryonic development. Cell 155, 1104–1118 (2013).

22. Zhang, B. et al. A human embryonic limb cell atlas resolved in space and time. Nature 1–11 (2023) doi:10.1038/s41586-023-06806-x.

23. Pickering, J. et al. An intrinsic cell cycle timer terminates limb bud outgrowth. eLife 7, e37429 (2018).

24. Logan, M. et al. Expression of Cre Recombinase in the developing mouse limb bud driven by a Prxl enhancer. Genesis 33, 77–80 (2002).

25. Currie, J. D. et al. The Prrx1 limb enhancer marks an adult subpopulation of injury-responsive dermal fibroblasts. Biology Open 8, (2019).

26. Markman, S. et al. A single-cell census of mouse limb development identifies complex spatiotemporal dynamics of skeleton formation. Developmental Cell 58, 565–581.e4 (2023).

27. Hara, K. & Ide, H. Msx1 expressing mesoderm is important for the apical ectodermal ridge (AER)-signal transfer in chick limb development. Dev Growth Differ 39, 705–714 (1997).

28. Subramanian, A. et al. Single cell census of human kidney organoids shows reproducibility and diminished off-target cells after transplantation. Nat Commun 10, 5462 (2019).

29. Krieg, M. et al. Tensile forces govern germ-layer organization in zebrafish. Nat Cell Biol 10, 429– 436 (2008).

30. Kosher, R. A., Savage, M. P. & Chan, S.-C. In vitro studies on the morphogenesis and differentiation of the mesoderm subjacent to the apical ectodermal ridge of the embryonic chick limb-bud. Journal of Embryology and Experimental Morphology 50, 75–97 (1979).

31. Akiyama, H., Chaboissier, M.-C., Martin, J. F., Schedl, A. & de Crombrugghe, B. The transcription factor Sox9 has essential roles in successive steps of the chondrocyte differentiation pathway and is required for expression of Sox5 and Sox6. Genes Dev 16, 2813–2828 (2002).

32. Firulli, B. A. et al. Altered Twist1 and Hand2 dimerization is associated with Saethre-Chotzen syndrome and limb abnormalities. Nat Genet 37, 373–381 (2005).

33. Gyllborg, D. et al. Hybridization-based in situ sequencing (HybISS) for spatially resolved transcriptomics in human and mouse brain tissue. Nucleic Acids Res 48, e112 (2020).

34. Manfrin, A. et al. Engineered signaling centers for the spatially controlled patterning of human pluripotent stem cells. Nat Methods 16, 640–648 (2019).

35. Li, P. et al. Morphogen gradient reconstitution reveals Hedgehog pathway design principles. Science 360, 543–548 (2018).

36. Boehm, B. et al. The Role of Spatially Controlled Cell Proliferation in Limb Bud Morphogenesis. PLoS Biol 8, e1000420 (2010).

37. Lázaro, J. et al. A stem cell zoo uncovers intracellular scaling of developmental tempo across mammals. Cell Stem Cell 30, 938–949.e7 (2023).

38. Masaki, H. & Ide, H. Regeneration potency of mouse limbs. Development, Growth & Differentiation 49, 89–98 (2007).

39. Aztekin, C. et al. Identification of a regeneration-organizing cell in the *Xenopus* tail. Science 364, 653–658 (2019).

40. Aztekin, C. et al. Secreted inhibitors drive the loss of regeneration competence in Xenopus limbs. Development 148, (2021).

41. Zhong, J. et al. Multi-species atlas resolves an axolotl limb development and regeneration paradox. Nat Commun 14, 6346 (2023).

42. Ran, F. A. et al. Genome engineering using the CRISPR-Cas9 system. Nat Protoc 8, 2281–2308 (2013).

43. Sybirna, A. et al. A critical role of PRDM14 in human primordial germ cell fate revealed by inducible degrons. Nat Commun 11, 1282 (2020).

44. Gritti, N. et al. MOrgAna: accessible quantitative analysis of organoids with machine learning. Development 148, dev199611 (2021).

45. Herrera, A., Saade, M., Menendez, A., Marti, E. & Pons, S. Sustained Wnt/β-catenin signalling causes neuroepithelial aberrations through the accumulation of aPKC at the apical pole. Nat Commun 5, 4168 (2014).

46. Choi, H. M. T. et al. Third-generation in situ hybridization chain reaction: multiplexed, quantitative, sensitive, versatile, robust. Development 145, (2018).

47. Hao, Y. et al. Integrated analysis of multimodal single-cell data. Cell 184, 3573–3587.e29 (2021).

48. González-Velasco, Ó., et al. Identifying Similar Astrocyte Populations in Motor Cortex and Spinal Cord Across Independent Single Cell Studies Without Data Integration. SSRN Scholarly Paper at 10.2139/ssrn.4765369 (2024).

49. Forster, B., Van De Ville, D., Berent, J., Sage, D. & Unser, M. Complex wavelets for extended depth-of-field: a new method for the fusion of multichannel microscopy images. Microsc Res Tech 65, 33–42 (2004).

50. Muhlich, J. L. et al. Stitching and registering highly multiplexed whole-slide images of tissues and tumors using ASHLAR. Bioinformatics 38, 4613–4621 (2022).

51. Patterson, H. & Manz, T. NHPatterson/wsireg: wsireg v0.3.5. Zenodo 10.5281/zenodo.6561996 (2022).

52. Mantes, A. D. et al. Spotiflow: accurate and efficient spot detection for imaging-based spatial transcriptomics with stereographic flow regression. 2024.02.01.578426 Preprint at 10.1101/2024.02.01.578426 (2024).

53. Axelrod, S. et al. starfish: scalable pipelines for image-based transcriptomics. Journal of Open Source Software 6, 2440 (2021).

54. Schmidt, U., Weigert, M., Broaddus, C. & Myers, G. Cell Detection with Star-Convex Polygons. in Medical Image Computing and Computer Assisted Intervention – MICCAI 2018 (eds. Frangi, A. F., Schnabel, J. A., Davatzikos, C., Alberola-López, C. & Fichtinger, G.) 265–273 (Springer International Publishing, Cham, 2018). doi:10.1007/978-3-030-00934-2_30.

55. Wolf, F. A., Angerer, P. & Theis, F. J. SCANPY: large-scale single-cell gene expression data analysis. Genome Biology 19, 15 (2018).

56. Allou, L. et al. Non-coding deletions identify Maenli lncRNA as a limb-specific En1 regulator. Nature 592, 93–98 (2021).

57. Desanlis, I., Paul, R. & Kmita, M. Transcriptional Trajectories in Mouse Limb Buds Reveal the Transition from Anterior-Posterior to Proximal-Distal Patterning at Early Limb Bud Stage. J Dev Biol 8, E31 (2020).

